# Environmental matrix and moisture are key determinants of microbial phenotypes expressed in a reduced complexity soil-analog

**DOI:** 10.1101/2024.10.02.616266

**Authors:** Josué Rodríguez-Ramos, Natalie Sadler, Elias K. Zegeye, Yuliya Farris, Samuel Purvine, Sneha Couvillion, William C. Nelson, Kirsten Hofmockel

**Affiliations:** Pacific Northwest National Laboratory, Earth and Biological Sciences Directorate

**Author notes:** Josué Rodríguez-Ramos and Natalie Sadler contributed equally to this work. Author order was determined based on who had the most time to address edits in final manuscript.

## Abstract

Soil moisture and porosity regulate microbial metabolism by influencing factors such as redox conditions, substrate availability, and soil connectivity. However, the inherent biological, chemical, and physical heterogeneity of soil complicates laboratory investigations into microbial phenotypes that mediate community metabolism. This difficulty arises from challenges in accurately representing the soil environment and in establishing a tractable microbial community that limits confounding variables. To address these challenges in our investigation of community metabolism, we use a reduced-complexity microbial consortium grown in a soil analog using a glass-bead matrix amended with chitin. Long-read and short-read metagenomes, metatranscriptomes, metaproteomes, and metabolomes were analyzed to test the effects of soil structure and moisture on chitin degradation. Our soil structure analog system greatly altered microbial expression profiles compared to the liquid-only incubations, emphasizing the importance of incorporating environmental parameters, like pores and surfaces, for understanding microbial phenotypes relevant to soil ecosystems. These changes were mainly driven by differences in overall expression of chitin-degrading *Streptomyces* species and stress-tolerant *Ensifer*. Our findings suggest that the success of *Ensifer* in a structured environment is likely related to its ability to repurpose carbon via the glyoxylate shunt while potentially using polyhydroxyalkanoate granules as a C source. We also identified traits like motility, stress resistance, and biofilm formation that underlie the degradation of chitin across our treatments and inform how they may ultimately alter carbon use efficiency. Together our results demonstrate that community functions like decomposition are sensitive to environmental conditions and more complex than the multi-enzyme pathways involved in depolymerization.

**Importance:** Soil moisture and porosity are critical mediators of microbial metabolism by influencing factors such as redox conditions, substrate availability, and soil connectivity. However, identifying how microbial community metabolism shifts in response to varying levels of moisture and porosity remains a challenging frontier. This difficulty arises from challenges in accurately representing the soil environment and in establishing tractable microbial communities that limit confounding variables. Moreover, inferring phenotypes based on “key” genes often fails to predict complex phenotypes that arise from cellular interactions. Here, we establish a tractably complex microbial community in a soil analog system amended with chitin and leverage it to understand how microorganisms respond to changes in porosity and moisture. By using genome-resolved metagenomics, metatranscriptomics, and metaproteomics, we report on the microbial lifestyle strategies that underpin changes in community expression like carbon conservation, biofilm production, and stress response.

## Introduction

To accurately understand the impact of environmental changes such as drought on terrestrial carbon cycling, it is crucial to understand their effects on soil microbiome function. However, heterogeneity in the biological, chemical, and physical properties of soil adds layers of complexity to understanding microbial organic matter (OM) decomposition. For example, soil particles and aggregates offer surface area and form a porous matrix with varying aqueous conditions ranging from saturated pore spaces to films too narrow for microbial movement (1). Further, porosity allows for compartmentalization and the privatization of goods (e.g., enzymes, osmolytes, vitamins) which translates to an expanded range of functional niches and increased diversity of organisms, functions, and metabolites, all of which in turn can alter interspecies competition (2). However, while the role of environmental matrices and moisture in shaping the soil microbiome is well acknowledged (3), mechanistic, genome-resolved details on how microbial communities alter their metabolism in response to these environmental changes are less studied. As such, there is a critical need for methods that enable scientists to interrogate changes at community and organismal levels.

Chitin is the key structural component for fungal cell walls, protists, and insect exoskeletons, and is the second most abundant biopolymer found on the planet, following cellulose (4, 5). Unlike cellulose, chitin is composed of carbon and nitrogen and, therefore, represents an essential elemental cycling hub (6). Like many polymers comprising soil OM, chitin is an insoluble substrate that requires enzymatic depolymerization into small subunits before microbial assimilation. As such, chitin decomposition rates are influenced by the effects that moisture can have on microbial metabolism. These metabolic shifts can occur because soil moisture regulates ecosystem characteristics like habitat connectivity, microbial mobility, and substrate diffusion (7–9) which ultimately influence changes in microbial strategies for yield (often measured as carbon use efficiency), resource acquisition (the production of extracellular enzymes, transporters, and motility), and/or stress response (e.g., oxygen stress, drought stress, etc.) (10). The chitin degradation phenotype requires the coordinated expression and deployment of a suite of enzymes for polymer hydrolysis (i.e., a resource acquisition). These enzymes include extracellular, cell tethered, or intracellular proteins such as AA10 (chitin monooxygenase), GH18/19 chitinase, and a range of enzymes involved in chitin oligomer/monomer transport and metabolism (11), ultimately making it a useful carbon source for identifying shifts in microbial lifestyle strategies. However, identifying how diverse metabolic strategies of microbial populations contribute to community decomposition in spatially heterogeneous environments remains a challenging frontier for soil ecology.

Extrapolating from an organism’s genetic code to its observable phenotypes is regarded as the “holy grail” of genomics research (12). However, microbial community phenotypes can often result from multiple complementary processes that can be difficult, and sometimes impossible, to measure accurately. These properties are described as emergent properties, meaning that they cannot be wholly predicted from the observations of the constituent components to the overall unit or process (13, 14), and instead are the result of a combination of factors. Examples of these emergent properties include substrate uptake involving multiple genes (14), and shifts in bacterial lifestyle strategies between high yield, resource acquisition, and stress tolerance in response to nutrient limitation or changing environmental conditions (10). Due to these emergent properties, inferring phenotypes based on the presence of specific key genes often fails to predict complex phenotypes that arise from cellular interactions like biofilm production and organismal metabolic handoffs, all of which can significantly alter community OM decomposition. Therefore, a comprehensive metabolic accounting beyond just “key” genes of specific processes is often required to fully understand microbial community dynamics.

Here, we present a mechanistic, genome-resolved overview of the impacts that the environmental matrix and moisture have on chitin decomposition by using a tractably complex microbial community. Specifically, we aimed to examine how the environmental matrix and moisture influence community structure, metabolism, and carbon use efficiency (CUE). To achieve our goal, we used a soil-derived microbial community (15) in either liquid culture or porous glass bead incubations at three different moisture levels. These moisture levels included 100%, 50% and 25% moisture saturation (**Fig. 1**).

**Figure 1:**
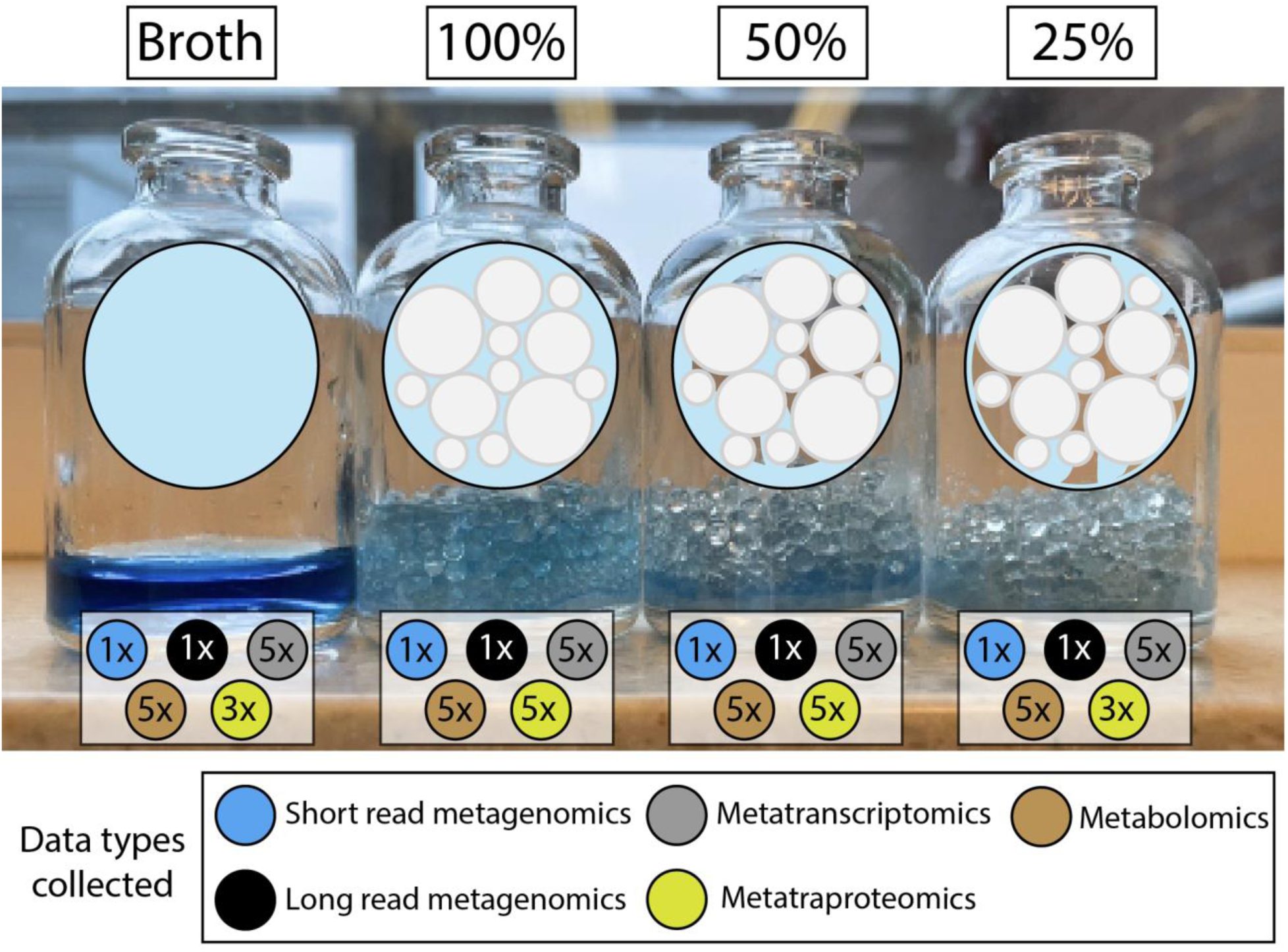
A multi-omic approach for understanding reduced complexity microbial community dynamics upon structure and moisture gradients. Overview of the experimental design that denotes the total number of samples that were collected per treatment type. The first bottle contains broth culture that is fully saturated, and the second bottle contains 100% saturation with porous media composed of glass beads (soil structure analog). The third and fourth bottles represent a decrease in moisture gradient. Circles below each bottle denote data types collected.

## Materials and Methods

### Soil extract preparation and characterization

The soil extract was prepared by mixing 500 g of sieved (2 mm sieve) soil from the Tall Wheatgrass Irrigation Field Trial in Prosser, WA with 1 L of Milli-Q water. The soil slurry was shaken at 160 RPM for 48 hr at 4°C. The slurry was then centrifuged 12,000xg for 30 min. Next, the supernatant (soil extract) was filter sterilized using a 0.22 µm vacuum filtration unit and stored at 4°C. The total C/N and elemental composition was determined by VarioTOC analyzer (Elementar, Langenselbold, Germany) for 5 replicates. The average total C was 48.52 µg/mL and average total N was 10.18 µg/mL for the soil extract.

### Cultivation and preparation of MSC-1 inoculum

MSC-1 inoculum was prepared by first cultivating the cells from glycerol stock on chitin-soil extract agar plates. To prepare chitin-soil extract agar plates for starting MSC-1 cultures, soil extract, 0.1% chitin (Bean Town Chemical), and 1.5% agar (BD Difco^TM^) were mixed and autoclaved 120°C for 15 min. For each 100 mm plate, 25 mL of chitin-soil extract agar solution was poured and allowed to cool at room temperature until solidified. Each plate was streaked with 200 µL of MSC-1 glycerol stock and incubated at 20°C in the dark for 7 days. On the 7^th^ day, the biomass was collected and transferred to a 50 mL conical tube. The cell pellets were suspended with 1X sterile PBS, centrifuged 7,000xg for 5 min, and the supernatant removed. Sterile soil extract enriched with 1000 ppm chitin and 600ppm glucose was then used to dilute the biomass to achieve an OD _600_nm of 0.09.

### Incubation preparation, incubation, and respiration measurements

We used 4 types of incubations for testing structure and moisture induced differences in community growth and phenotypes: liquid culture, 20%, 10%, and 5% w/w which translates to liquid culture, 100%, 50% and 25% moisture saturation. To test structure, we prepared standard unstructured liquid incubations by adding 3 mL of sterile water and 1 mL of soil extract carrying MSC-1 inoculum prepared as described above to sterile 30 mL serum bottles. The same volume of water and soil extract inoculum was added to 30 mL serum bottles with a porous matrix composed of 16 g of mixed size glass beads (2.7 mm diameter beads (10 g), 1.00 mm diameter beads (4 g), and 0.1mm diameter beads (2 g)). In addition to the glass beads, 0.1 g of 45-90 µm polyacrylamide beads (Bio-Gel P-6, BioRad) were added to help uniformly distribute the moisture throughout the porous media. The 4 mL of liquid volume added to these incubations was 20% (w/w) liquid to glass beads and represents a 100% fully moisture saturated condition. To test varying moisture levels, we also prepared 50% and 25% moisture saturated incubations. The 50% and 25% saturated incubations had equivalent amounts of glass beads, polyacrylamide beads, and soil extract MSC-1 inoculum as the 100%, but the 50% received only 1 mL of sterile water and the 25% received no addition water (**Fig. 1**). Prior to inoculation, the serum bottles and beads were heat sterilized (121°C for 45 min), cooled to room temperature, and inoculated with 1 mL of soil extract liquid enriched with 1000 ppm chitin, 600ppm glucose, and MSC-1 cells adjusted to an OD_600_ nm of 0.09. In total, each sample received 722.92 µg of C (434.4 µg from chitin, 240 µg from glucose, and 48.53 µg from soil extract) and 73.48 µg of N (63.3 µg from chitin and 10.18 µg from soil extract) yielding a C:N ratio of 9.84:1. Next, sterile water was added as previously described. To acquire enough biomass for downstream analysis, we prepared 100 incubation bottles for each treatment and pooled 20 incubations for each of 5 composited samples. Culture bottles were weighed and then incubated at 28°C for 4 days. An additional 10 replicates for each treatment were also incubated for 8 days to continue monitoring respiration. Respiration for five representative replicates from each treatment were measured daily. To do so, 10 mL of head space gas was collected from incubations by 20 mL gastight syringe. The headspace gas was then injected in to and measured by EGM-4 PP infrared CO_2_ analyzer. Incubations were transferred to a sterile biological safety cabinet and septa were removed to allow for headspace gas exchange for 30min. Incubation bottles were then re-capped with septum and aluminum collar.

### Sample harvesting, subsampling, and preparation

After 4 days of incubation, samples were harvested for analysis. To generate enough biomass for metagenomic sequencing, 20 replicate incubations were pooled per composite sample to create 4 representative composite samples for each treatment. The liquid content of each sample replicate bottle was transferred to 50 mL conical collection tubes. Next, 2 mL of MES buffer (pH 6) was added to the bead matrix and vortexed for 10 sec. to suspend cells. The cell suspension was transferred to the corresponding collection tubes and this step was repeated once more with an additional 2 mL of MES buffer. The harvested and pooled samples were split into two aliquots. The first was comprised of 85% of the pooled sample volume which was used for downstream proteomic sample processing, while the remaining 15% of the sample volume was used for DNA/RNA extraction and extracellular metabolomic sample processing. The 85% and 15% volume sample aliquots were centrifuged at 5,000 x g for 10 min to pellet cells. The 15% volume cell pellets and supernatants were separated, and flash frozen with liquid N_2_ for nucleotide and metabolite extractions. The 85% volume samples were kept on ice or at 4C for additional processing.

The supernatant from the 85% samples were concentrated and buffer exchanged with fresh MES buffer using 30 kDa molecular weight cutoff centrifugation filters with PES membranes (Millipore). The washed extracellular proteins were then resuspended in 0.5 mL MES buffer and transferred to 1.5 mL conical tubes. Meanwhile, the 85% sample cell pellets were transferred with 0.5 mL of MES buffer to 2 mL mechanically resistant tubes holding 100 µL of zirconium beads (100 µm diameter, Biospec). The cells were then lysed by bead beating (1min at 6m/s, rest 5min, 3x). The 85% volume sample lysates and concentrated supernatants were then combined, and protein was quantified by CBQCA protein quantification kit assay (ThermoFisher) as described by the manufacturer with each sample run in triplicate using 5 µL per technical replicate. Protein was then quantified by Biotek Synergy H1 plate reader with 465nm excitation and 550nm emission. The supernatants from the 15% volume collection were thawed on ice and 1.5 mL were filtered using 0.22 µm syringe filters (Pall Corps) and transferred to 1.5 mL tubes. The samples were then covered with BreatheEasier membranes, flash frozen on liquid N_2_, and then lyophilized till dry.

### Metagenome extraction, assembly, and data processing

The cell pellets from the 15% volume collection were ZymoBIOMICS DNA/RNA mini prep kit was used for co-extracting both pools of nucleotides. RNA/DNA were quantified by Qubit assay and purity was checked by NanoDrop (absorbance ratios of 260nm/280nm and 260nm/230nm). Samples were shipped on dry ice to be analyzed. Both short-read and long-read sequences were processed at the Joint Genome Institute (JGI) with Illumina and PacBio CCS sequencers, respectively. Long-read metagenomes were fully assembled and processed using the JGI workflow (16) and downloaded from the JGI Data Portal. Interleaved, short-read sequences were downloaded from the JGI Data Portal and trimmed with bbduk (17) using parameters ktrim=r k=23 mink=11 hdist=1 qtrim=rl trimq=20 minlength=75. Trimmed reads were subsequently assembled with MEGAHIT (18) using default settings. Ribosomal contamination was removed by identifying ribosomal RNA using barrnap (19) and removing reads that mapped to an rRNA reference database.

### Bacterial metagenomic binning, quality control, annotation, and taxonomy

Metagenome assembled genomes (MAGs) were generated using both the standard JGI workflow (long-read sequencing) (16) and metaWRAP (short-read sequencing) (20). For short-read metagenomes, scaffolds ≥2500bp were extracted using seqkit (21) and binned using the MetaWrap binning and refinement modules with default settings. After binning, MAGs from both short-read and long-read metagenomes were pooled and processed together. To ensure only quality MAGs, any MAGs that were not medium quality (MQ) or high quality (HQ) per MIMAG standards (22) were discarded from subsequent analyses resulting in 174 MAGs. These MAGs were subsequently dereplicated at 99% identity yielding in 29 MAGs. MAG and sample statistics can be found in **Table S1**. These MAGs were used as the mapping reference for multi-omic recruitment as shown in **Fig. 2**.

**Figure 2:**
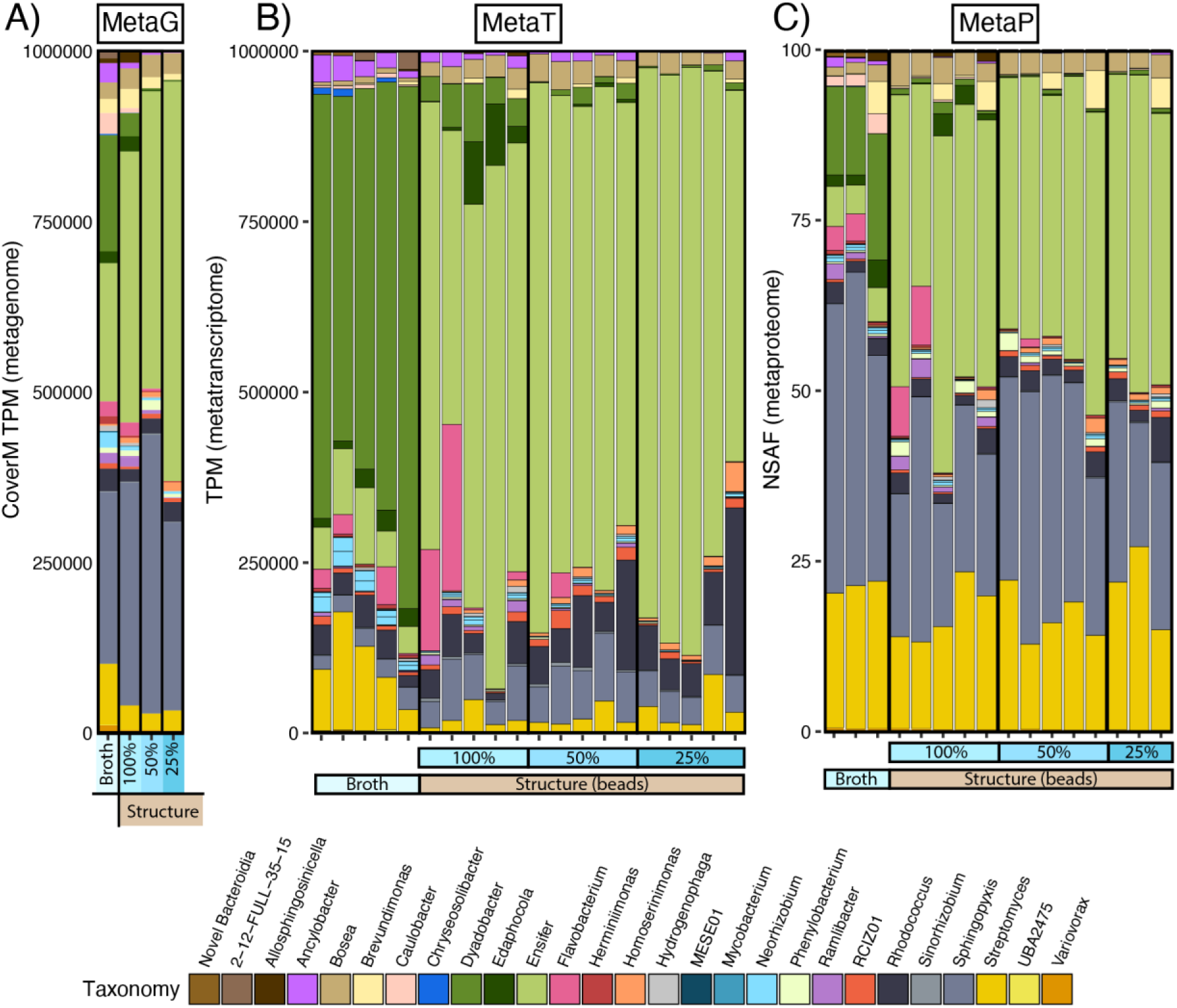
Multiple ‘omics data types highlight different expression patterns across experimental treatments. Stacked bar charts are colored by genera and denote normalized expression (see Materials and Methods) detected by either **A)** CoverM normalized metagenomics (n = 5), **B)** TPM normalized metatranscriptomics (n = 20) or **C)** NSAF normalized metaproteomics (n = 16).

These 29 MAGs were functionally annotated using DRAM (23) and eggNOG (24), with full annotations hosted on GitHub (https://github.com/jrr-microbio/MSC1_SSA_incubations_moisture), and summarized in the **Supplemental Results**, **Table S2** and **Table S3**. Aside from DRAM-assigned cutoffs for the genomic potential of metabolisms, we employed additional cutoffs for assigning potential in our functional dot plot in the supplemental results. Overall, for any respiratory or fixation metabolisms to be assigned, genomes must encode at least 50% of all genes in each pathway. In addition to that base cut-off: for reductive TCA cycle (i.e., Arnon-Buchanan cycle) genomes had to encode Citrate lyase (K15230, K15231), citryl-CoA lyase (K15234), or citryl-CoA synthetase (K15232, K15233) (25); for reductive pentose phosphate pathway (Calvin-Benson-Bassham cycle) genomes had to encode the rate limiting ribulose-1-5 bi-phosphate carboxylase / oxygenase (Rubisco) gene (K01601, K01602) (26); for 3-Hydoxypropionate bi-cycle and Dicarboxylate-4-hydroxybutyrate cycle genomes had to encode 4-Hydroxybutyrate dehydrogenase (K00043, K08318, K18120). For complex carbon degradation and CAZYmes, DRAM calls were kept with the addition that chitin degradation genomes must contain GH18 or GH19. All GH18 and GH19 chitinases had to be confirmed by AlphaFold to be structurally like known chitinases and must have either: 1) a reverse best hit to KEGG chitinase, or 2) a consensus of 2 different annotation metrics (i.e., KEGG, PFAM, CAZy, eggNOG). Chitin degradation is outlined in **Fig. 3**. All genes involved for the glyoxylate shunt are described in **Fig. 4**. For ectoine synthesis functional call, genomes had to encode ectoine synthase (K06720), for ectoine synthesis bar charts in **Fig. 5**, genes included in query were: K00928, K12524, K12525, K12526, K00836, K06718, K10674, K06720, K03735, K03736 with pathway described in (27). For ectoine degradation functional call, genomes had to encode ectoine hydrolase (K15783) with pathway described in (27). For N-acetylputrescine synthesis bar charts in **Fig. 5**, genes included in query were: K11073, K11076, K11075, K11074, K01585, K10536, K12251, K01581, K01480 with pathway described in (28). Raw DRAM metabolism summary output is on Github (https://github.com/jrr-microbio/MSC1_SSA_incubations_moisture). For taxonomic classification, we used the Genome Taxonomy Database (GTDB) toolkit version 2.3.2 using the r214 database on October, 2023 (29, 30). PSORTb web-based GUI was used to localize chitinase enzymes of interest (31). All supplementary code used to generate these data are hosted on GitHub (https://github.com/jrr-microbio/MSC1_SSA_incubations_moisture).

**Figure 3:**
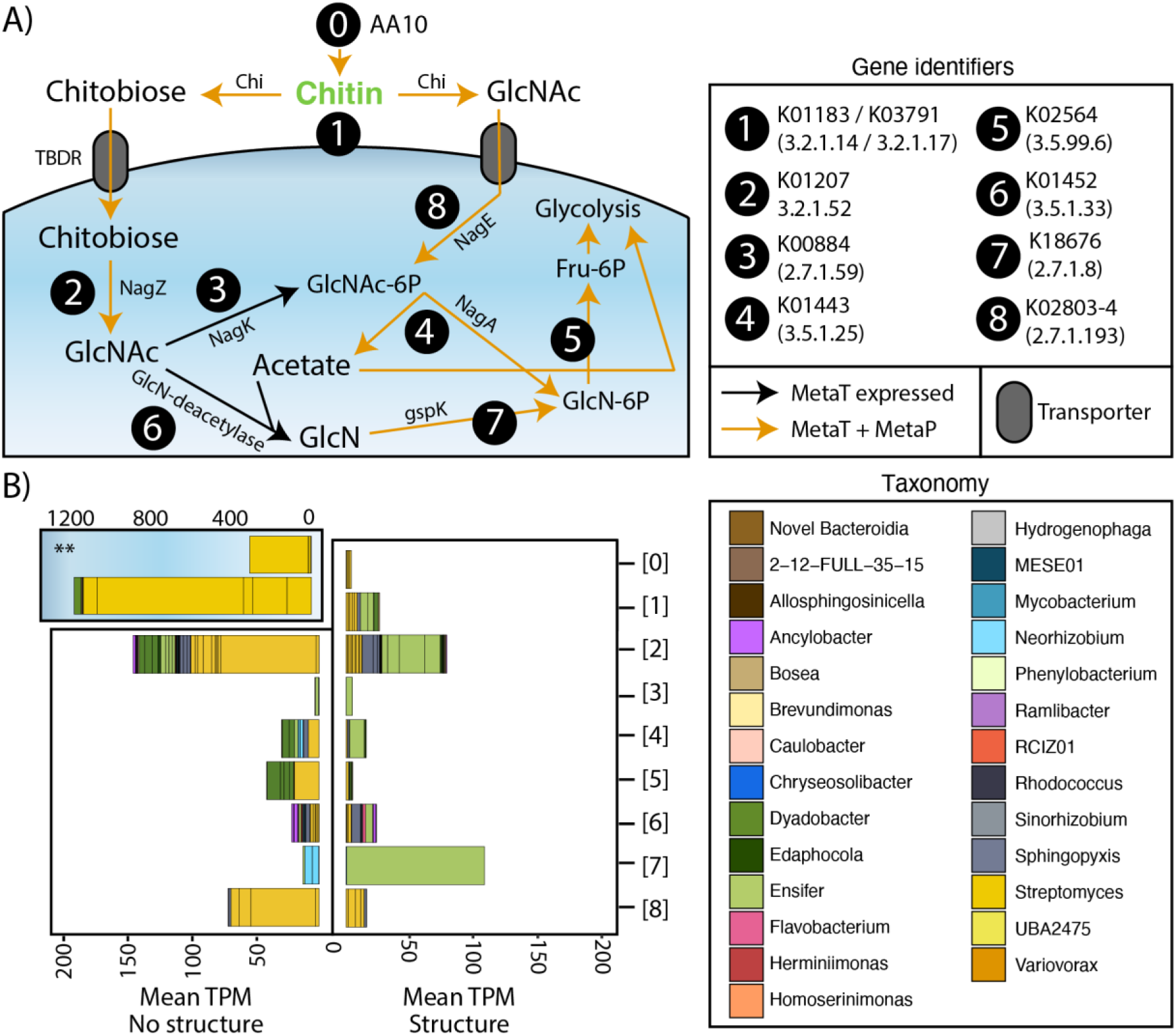
Community members express genes related to chitin degradation. **A)** Schematic diagram of a cell shows the overall pathways of chitin degradation in a generic cell. Each gene is represented by a circle and labeled by their KO, common name, and E.C. number. Arrows denote detection in metatranscriptome (black) or both metatranscriptome and metaproteome (orange). Butterfly plots denote the overall mean TPM of each of the genes denoted in (A) for either liquid culture (left) or soil analog 100% moisture (right). Colors on stacked bars show genera-level GTDB taxonomy. TBDR = TonB dependent transporter. **Note: Axis for “0” (AA10) and “1” (GH18 / GH19) is different.

**Figure 4:**
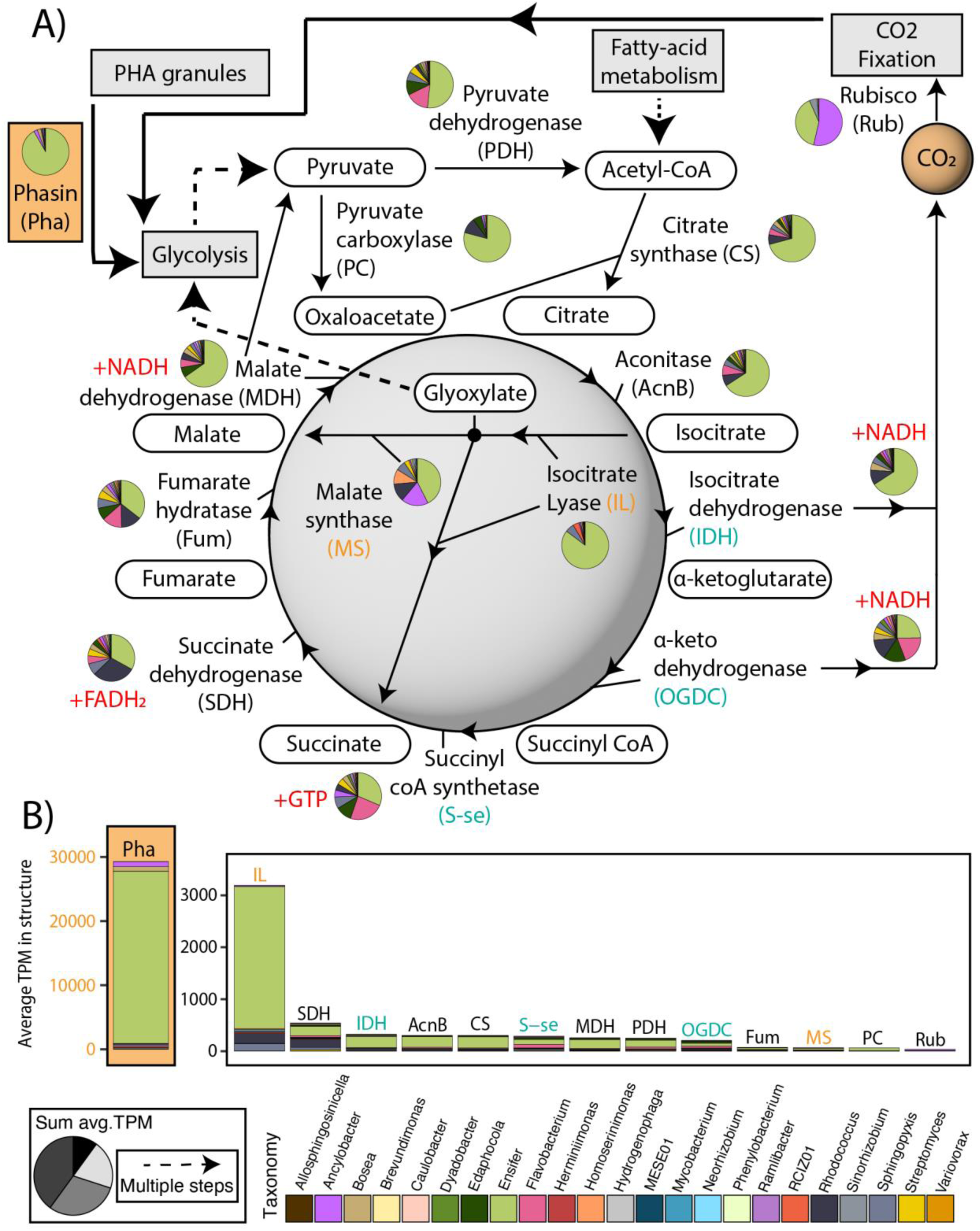
Community-wide transcriptional response in favor of the glyoxylate shunt. **A)** Diagram shows the TCA cycle overlaid with the glyoxylate shunt. Pie charts denote the proportion of the summed averages for the structure treatment of each of the genes and are colored by the genera of the organism expressing the gene. **B)** Bar charts denote the summed average TPM for the structure treatment of all the genes within each organism that contribute to each unique step in **(A)**.

**Figure 5:**
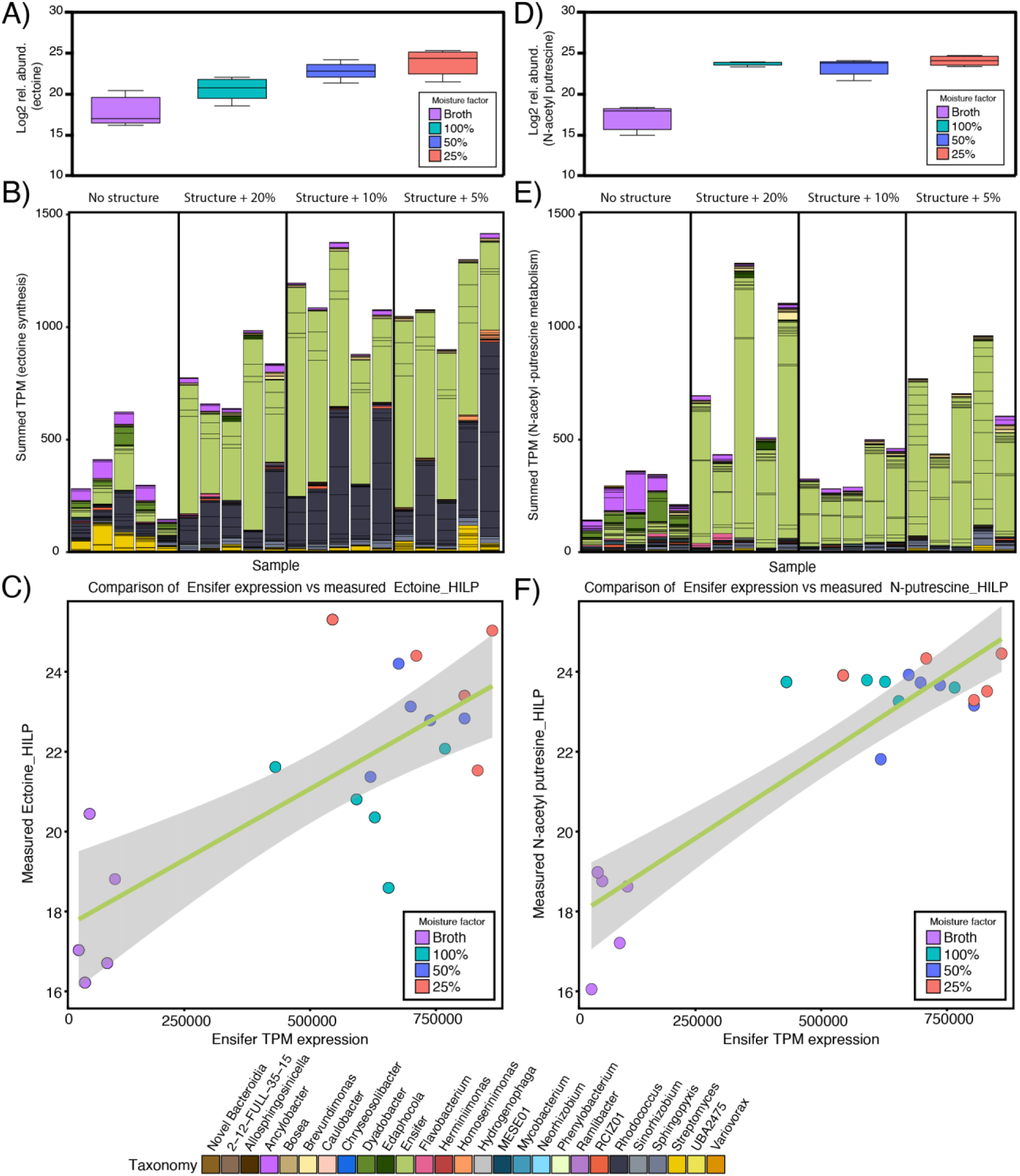
Ensifer is key player upon introduction structure and moisture likely due to its ability to respond to stress and possibly form biofilms. A) Bar plots show the log-2-fold change of significantly different (p ≤ 0.05) ectoine metabolite. B) Stacked bar chart shows the summed TPM expression of ectoine synthesis genes colored by phyla of contributing organisms. C) sPLS shows the measured ectoine concentrations versus the overall Ensifer TPM expression. D-F) Same as A-C but for significantly different (p ≤ 0.05) N-acetyl putrescine metabolite.

### Genome relative abundances (metagenomics) and relative expression (metatranscriptomics)

To determine the relative abundance and expression of our MAGs, metagenomic and metatranscriptomic datasets were mapped to a reference database of the 29 MQ/HQ MAGs using bbmap from the bbtools suite (17). To remove ribosomal RNA (rRNA) contamination from metatranscriptomes, reads were mapped to a database of rRNA from our assembled metagenomes identified using barrnap (19) with flags ambiguous=all and perfectmode=t. Reads that perfectly mapped to rRNA were then removed and subsequent cleaned reads were used. Both cleaned metatranscriptomic reads and metagenomic reads were mapped to our MAG database using bbmap with flag ambiguous=all. After initial mapping, SAM files were then filtered to report hits of ≥98% identity (minidfilter=0.98), and only primary mappings (pimaryonly=t) using reformat.sh from bbsuite (17). Metagenomic reads were additionally filtered to include only paired mapping results (paired=true). Metagenomic mapping SAM files were processed with coverM (32) and were normalized using the TPM calculation (for gene and library size), a minimum contig coverage of 75%, and a minimum depth coverage of 2x. Metatranscriptomic mapping SAM files were processed with featureCounts (33) using the subread package from Bioconda (34). Genes of the 29 MAG database were called using DRAM (23), and the resulting .gff file was used for featureCounts along with flags -t rrna, trna, CDS, -s 2 (reversely stranded), -M (multi-map), and -p (paired). The resulting counts were then normalized in R to the total length of each respective gene and subsequently converted to transcripts per million (TPM) according to the code in Additional File 4 of the geTMM manuscript (35).

Raw counts for the read mapping that met minimum contig coverage cutoffs were used for DESEQ2 (36), with adj. p values ≤ 0.05 reported (**Table S4**). For NMDS of metatranscriptomic data, normalized TPM values for each MAG were used as input in R for metaMDS from the Vegan library (37). Normalized TPM values for each MAG were also used for sPLS (38). All R plots aside from **Fig. S5** (which was made using RawGraphs (39) were made in R using ggplot2 (40). All supplementary code used to generate these data are included hosted on GitHub (https://github.com/jrr-microbio/MSC1_SSA_incubations_moisture). Read mapping results can be found in **Table S1** and DESEQ2 raw results are in **Table S4**.

### Metaproteome generation and peptide mapping

Aliquots of 100 µg of each protein sample were denatured with 8M urea and reduced by incubating at 60 °C for 30 min with 500 mM dithiothreitol. Samples were then alkylated by incubating with 40 mM at 37 °C for 60 min. Digestion was performed using 1:50 ratio of trypsin (Promega, Madison, WI, USA) to protein and incubated for 3 h at 37 °C. Digested peptides were loaded on C18 SPE columns, preconditioned with MeOH and equilibrated with 0.1% trifluoroacetic acid (TFA). Samples were washed with a solution of 5% acetonitrile (ACN) and 0.1% TFA and eluted with a solution of 20% ACN and 0.1% TFA. Eluted peptides were dried by speed vacuum and reconstituted in 25 mM NH_4_HCO_3_ (pH 8), and the concentrations were normalized to 0.1 g L^-1^ for analysis.

Peptides were analyzed using a custom-built reversed-phase LC column coupled to a LTQ Orbitrap Velos mass spectrometer (Thermo Fisher). The raw spectra from the MS were extracted using MSConvert (part of the Proteowizard tool suite (41) and precursor m/z values re-calibrated using MZRefinery (42) and searched against the genes of all long-read and short-read metagenomes using MS-GF+ (41) in target-decoy mode (43). MS-GF+ Q-Values were used to calculate the FDR value (∼0.01 cut off). Peptide-to-protein relationships were established using parsimony such that only non-unique peptides belonging to more than one uniquely identified protein were not used for further quantitation.

Detected metaproteome peptides were then mapped back to the genes from our total metagenomic assemblies reporting all possible hits. Results from the peptide matching of total metagenomic gene content were then subset to only those genes that were present within our 29 MAG database when peptides were confidently assigned to one or more genes, and subsequent analyses were performed on this subset. Metaproteomic counts were subsequently converted to spectral abundance factors (SAF) by calculating the length of each protein and normalizing by total counts per sample (NSAF). For NMDS of metaproteomic data, normalized NSAF values for each MAG were used as input in R for metaMDS from the Vegan library (37). Raw metaproteomic counts as well as the NSAF normalized values for the database of 29 MAGs are shown in **Table S1**.

### Extracellular LC-MS/MS metabolite processing

Extracellular metabolites were analyzed using reverse phase (RP) and hydrophilic interaction chromatography (HILIC) separations on a Thermo Fisher Scientific Q Exactive Plus mass spectrometer (Thermo Scientific, San Jose, CA) coupled with a Waters Acquity UPLC H class liquid chromatography system (Waters Corp., Milford, MA). Metabolites were brought up in 100 µL of 80% LCMS grade methanol and 20% nanopure water, vortexed to mix, and centrifuged 4,500 rpm for 5 minutes. The wells of a 0.2 µm PTFE 96 well filter plate were preconditioned with methanol and then placed on top of a 96 well collection plate. Samples were then transferred to the filter plate wells and N_2_ was applied at 9 psi via a CEREX System 96 processor for positive pressure SPE to push the sample through the filtered well plate into the collection well plate.

RP separation was performed by injecting 5 µL of sample onto a Thermo Scientific Waters Acquity UPLC BEH C18 column (130 Å, 1.7 µm, 2.1 mm ID X 100 mm L) preceded by a Acquity UPLC BEH C18 Vanguard Pre-Column (130 Å, 1.7 µm, 2.1 mm ID X 5 mm L) heated to 40°C. Metabolites were separated using a 15-minute gradient with data collected on the first 10 minutes. For RP acquisition, positive mode polarity was run. The gradient used was identical, but the solvent composition was different between the modes. The positive mode mobile phase A consisted of 0.1% formic acid in nanopure water with the mobile phase B consisting of 0.1% formic acid in LCMS grade methanol, while the negative mode mobile phase A consisted of 6.5 mM ammonium bicarbonate in nanopure water at a pH of 8 with the mobile phase B consisting of 6.5 mM ammonium bicarbonate in 95% LCMS grade methanol and 5% nanopure water. The gradient used was as follows (min, flowrate in mL/min, %B): 0,0.35,5; 4,0.35,70; 4.5,0.35,98; 5.4,0.35,98; 5.6,0.35,0.5; 15,0.35,0.5. HILIC separation was performed by injecting 3 µL of sample onto a Waters Acquity UPLC BEH Amide column (130 Å, 1.7 µm, 2.1 mm ID X 100 mm L) preceded by a Acquity UPLC BEH Amide Vanguard Pre-Column (130 Å, 1.7 µm, 2.1 mm ID X 5 mm L) heated to 40°C. Metabolites were separated using a 21-minute gradient with data collected for the first 16 minutes. For HILIC mode, both positive and negative spray ionizations were used in separate injections using the same mobile phase compositions. The HILIC mobile phase A used consisted of 0.125% formic acid and 10 mM ammonium formate in nanopure water with a mobile phase B consisting of 0.125% formic acid and 10mM ammonium formate in 95% LCMS grade acetonitrile and 5% nanopure water. The gradient used was as follows (min, flowrate in mL/min, %B): 0,0.4,100; 2,0.4,100; 5,0.4,70; 5.7,0.4,70; 7,0.4,40; 7.5,0.4,40; 8.25,0.4,30; 10.75,0.4,100. For both RP and HILIC separations the Thermo Fisher Scientific Q Exactive was equipped with a HESI source and high flow needle with the following parameters: spray voltage of 3.6 kV in positive mode and 3 kV in negative mode, capillary temperature of 350 °C in positive mode and 275 °C in negative mode, aux gas heater temp of 325 °C in positive mode and 300 °C in negative mode, sheath gas at 45 L/min in positive mode and 30 L/min in negative mode, auxiliary gas at 15 L/min in positive mode and 25 L/min in negative mode, and spare gas at 1L/min in positive mode and 2 L/min in negative mode. Metabolites were analyzed at a resolution of 70 k and a scan range of 70 to 1000 m/z for parent ions followed by MS/MS HCD fragmentation which is data dependent on the top 4 ions with a resolution of 17.5 K and stepped normalized collision energies of 20, 30, and 40. All metabolite data is available in **Table S5**.

#### Metabolomics data analysis

Metabolite identifications were made using MS-DIAL (44) v4.92 for peak detection, identification and alignment. The experimental data was matched both to in-house libraries (m/z less than 0.003 Da, retention time less than 0.3 min, MS/MS spectral match) and a compilation of publicly available MS/MS databases (m/z less than 0.003 Da, MS/MS spectral match) available in MS-DIAL. The tandem mass spectra and corresponding fragment ions, mass measurement error, and aligned chromatographic features were manually examined to remove false positives. Relative quantification was performed by calculating peak areas on extracted ion chromatograms of precursor masses. Features detected in at least three of the five replicates were retained.

### Statistical analyses

All statistical analyses were performed with the TPM normalized or NSAF normalized values described above. To determine the overall composition of our microbial community across treatments, we calculated Bray-Curtis dissimilarities using vegan (37) in R, and these were visualized using Nonmetric multidimensional scaling (NMDS) with k=2. For each treatment type, MANOVA analyses were then performed using the adonis2 function from vegan. To identify differentially expressed genes, raw counts that met the 75% alignment fraction and 2x coverage were input into DESeq2 in R (36). Samples were split into two treatments: structure comparisons and moisture comparisons. The subset of moisture comparison samples that were compared were those corresponding to 5% and 25% moisture. DESeq2 was then run on each subset of samples separately. To identify which genomes were strongly related to the collected metabolites, we used sparse partial least square regressions (sPLS) (38) on the metatranscriptomic expression patterns of the structure and moisture sample subsets. For metabolism-level differential expression, all annotations and calls that were used to aggregate groups are denoted in **Table S2** and **Table S6**, and detailed inputs are on GitHub https://github.com/jrr-microbio/MSC1_SSA_incubations_moisture.

Statistical analysis of metabolomics data was performed using the PMart web application (45). Data was log2-transformed and normalized via global median centering. Statistical comparisons were performed for structure (broth vs 100% moisture) or moisture (100%, 50%, or 25% moisture) using ANOVA with a Holm test correction (46). For each metabolite, these adjusted p values and the mean log2 fold changes for each of the above comparisons are reported (**Table S5**), along with the number of observations per group.

## Results

### Leveraging multi-omics datasets of a reduced complexity microbial community to understand the role of environmental matrix and moisture on community patterns

After a 96-hr incubation of our soil-derived microbial community (15) in either liquid culture or soil analogs, we sequenced 4 long-read reference metagenomes, 4 short-read reference metagenomes, 20 metatranscriptomes, and 16 metaproteomes (**Fig. 1**). From our metagenomes, we generated a reference database of 29 metagenome assembled genomes (MAGs) that were medium-quality or greater (average completion = 94.0%, average contamination = 0.4%). In fact, 19 of our MAGs (66%) were from long-read metagenomes that had ≤ 3 scaffolds and were 98% complete. Among our MAGs were 6 of the 8 isolates that comprise the MSC-2 reduced complexity consortia (15), which have been extensively characterized. In total, our MAG database contained 128,133 genes.

To determine if microbial community expression was altered by our treatments, we recruited metatranscriptomic reads and metaproteomic spectra to our reference database of 29 MAGs (**Fig. 2**, **Table S1**) and normalized the abundances to total transcripts per million (TPM) and normalized spectral abundance factors (NSAF), respectively. On average, 60% of metatranscriptomic reads mapped back to the reference database. Metatranscriptome reads were recruited to 95,756 genes (75% of total), while metaproteomes had spectral hits to 8,627 genes (7% of total). Due to low recruitment in our metaproteomics data, we performed our statistical and metabolic analyses using transcriptomic data.

At a community level, overall expression by both metatranscriptomics and metaproteomics showed significant differences between soil analog and broth treatments (metaT: R^2^ = 0.86, p ≤ 0.05, metaP: R^2^ = 0.76, p ≤ 0.05) (**Fig. S1**). Soil analog moisture content (100%, 50%, 25% saturation) also showed a significant effect on community expression (metaT: R^2^ = 0.26, p ≤ 0.05), albeit at a less notable level than the liquid-to-soil analog comparisons. No effect of moisture content was detected in the metaproteome profiles, likely because of data sparsity (metaP: R^2^ = 0.09, p > 0.05). While moisture patterns explained less of the variation than the environmental matrix, our results showed that both habitat structure and moisture are strong determinants of microbiome community expression and highlighted that communities growing under broth conditions differ from those growing in structured habitats.

To explain which organisms were causing these community shifts, we interrogated the abundance and expression of individual MAGs across treatments (**Fig. 2**). In the liquid culture, *Dyadobacter* accounted for 62% of metatranscriptomic expression (and third highest metagenomic abundance), followed by *Streptomyces* (10% of expression, fourth highest metagenomic abundance) and *Ensifer* (7% of expression, second highest metagenomic abundance). In the soil analog communities, incubations with highest moisture (i.e., 100% saturation) showed an expression profile of 62% *Ensifer* (highest metagenomic abundance), 8% *Flavobacterium* (eighth highest metagenomic abundance), and 6% *Sphingopyxis* (second highest metagenomic abundance). Across the moisture treatments in the soil analogs, community abundance and expression were more similar. Specifically, in the soil analog community with lowest moisture (25% saturation), the community expression was 75% *Ensifer* (highest metagenomic abundance) followed by *Rhodococcus* (10% of expression, fifth highest metagenomic abundance) and *Sphingopyxis* (5% of expression, second highest metagenomic abundance). Throughout our incubations, some low abundance members disproportionally contributed to the overall expression pools (i.e., *Streptomyces*, *Dyadobacter*).

### Soil analog conditions lead to significantly lower chitin metabolism by Streptomyces

Changes in chitin decomposition phenotypes were detected via metatranscriptomic activity across our broth and soil analog treatments (**Fig. S1)**. Specifically, we detected high transcriptomic abundance of chitin degradation genes in the broth treatment, with *Streptomyces* GH18 chitinase being the second most highly expressed carbon degradation gene. In broth, 89% of chitin metabolism gene expression was assigned to *Streptomyces,* consistent with a central role in resource acquisition (**Fig. 3, Table S3**). While the capability to degrade chitin and its byproducts is widespread across our MAGs (**Fig. 3**), the other top 4 chitin degrading genera in broth (i.e., *Dyadobacter, Neorhizobium, Ensifer*, *Sphingopyxis*) together account for only 9% of total chitin degradation expression, and the remaining 22 genera account for less than 1%. While our metaproteomics dataset was sparse, it supported the observed transcriptomic patterns and showed 77% of the expressed peptides for chitin degradation were from *Streptomyces* in the broth, followed by *Dyadobacter* (12%) and Sphingopyxis (11%). These results support that *Streptomyces* is the key contributor to the decomposition of the chitin backbone in this reduced complexity broth system.

In the 100% soil analog incubations, the abundance of transcripts related to chitin degradation was significantly lower than in the broth (p ≤ 0.05). Aside from chitinases, the genes responsible for the linearization of chitin (AA10, lytic polysaccharide monooxygenase) had a 200-fold lower expression in 100% soil analog than in liquid culture. *Streptomyces* was the only organism that encoded and transcribed AA10 genes. An extracellular GH18 encoded by *Streptomyces,* which was the most highly transcribed DRAM-annotated gene in broth incubations, showed significantly lower expression in the 100% soil analog (base mean = 12,272, log_2_ fold change = -8.8, p ≤ 0.05). Further, all *Streptomyces*-encoded chitinases had significantly lower transcriptomic abundance in 100% soil analog (avg. log_2_ fold change = -6.5). This suggests that in a structured environment, the role of *Streptomyces* in chitin degradation is reduced.

After 4 days, we were unable to detect any expressed genes for the degradation of chitin in the soil analog incubations at the same expression level as in the liquid culture. In fact, chitin degradation only accounted for 0.9% of the total transcripts in soil analog conditions. In these soil analogs, *Ensifer* took over the largest proportion (69%) of chitin metabolism in the 100% moisture level, followed by *Streptomyces*, *Sphingopyxis*, *Dyadobacter,* and *Bosea* which account for an additional 25% (**Table S3)**. The remaining 22 MAGs provided only 6% of chitin-related transcriptomic abundance. However, contrary to the metatranscriptomic data, 61% of the measured peptides in the metaproteomes of the soil analog still belonged to *Streptomyces,* with *Ensifer* encoding 31%, and the remaining 9% coming from *Sphingopyxis*, suggesting that *Streptomyces* is likely still decomposing chitin in soil analog. Given that our communities were compared at day 4 after initial incubation, this may reflect temporal differences in the stability of RNA transcripts and proteins. Nonetheless, our metatranscriptomic and metaproteomic data suggest that microbial communities are more actively using chitin as a carbon source in liquid culture than in the 100% soil analog, and that *Streptomyces* is the major contributor to chitin backbone degradation in liquid.

### Coupled respiration measurements and metatranscriptomics suggest that the glyoxylate shunt likely sustains Ensifer cellular carbon needs

Since the overall expression of chitin metabolism genes only comprised a fraction of transcriptomic activity in the soil analog, we wanted to understand what other metabolic processes were supplementing microbial community carbon needs. Isocitrate lyase (*aceA* [K01637]) from *Ensifer* was observed to be highly expressed and had significantly higher expression in structured compared to broth incubations (base mean = 39,980, log_2_ fold change = 5.57, p ≤ 0.05) (**Fig. S2**). Isocitrate lyase is the first step in the glyoxylate shunt, a carbon conservation mechanism where isocitrate is transformed into glyoxylate and succinate, bypassing the carbon dioxide-generating portion of the TCA cycle (**Fig. 4A**). Highlighting the overall importance of this metabolism, while the response towards the use of glyoxylate and carbon conservation is mostly expressed by *Ensifer* possibly due to its overwhelming abundance, 19 of the other organisms in MSC-1 (i.e., *Rhodococcus* and *Sinorhizobium*) also encoded isocitrate lyase genes. Further, 3 of those (*Sinorhizobium, Ancylobacter and Phenylobacterium)* had significantly elevated levels of expression in soil analog 100%, albeit at possibly negligible levels (avg. base mean = 220, avg. log_2_ fold change = 6.76, p ≤ 0.05). In 100% soil analog, *Ensifer* isocitrate lyase showed 6-fold higher transcriptomic abundance than any other gene in the TCA cycle (**Fig. 4B**). *Ensifer* also expressed genes necessary to use the produced glyoxylate to generate 2-Phospho-D-glycerate (*gcl* [K01608], *hgy*/*gip* [K01816], *gyaR*/*GOR1* [K00015], and glxK/garK [K00865]), which can ultimately feed into pyruvate for glycolysis. Together, these data suggest that carbon conservation mechanisms via the glyoxylate shunt are likely supplementing the growth of our microbial consortia in soil analog.

Given that in soil analog 1) isocitrate lyase expression suggested carbon was being shunted and conserved, and 2) chitin degradation was lowly expressed, we wanted to understand the source of the carbon sustaining this community. The ability to store carbon as granules was encoded by 62% of the MSC-1 organisms. *Ensifer-*encoded phasins (e.g., the granule-associated proteins of polyhydroxyalkanoates) were the second and third most expressed genes in our entire dataset by metatranscriptomics. The two *Ensifer* phasins were also the most expressed genes with significantly higher transcript abundance in structured conditions when compared to liquid culture (avg. base mean = 65,134, log_2_ fold change = 4.25, p ≤ 0.05). These phasins had 67-fold higher expression than the isocitrate lyase from *Ensifer* and 380-fold higher than any TCA cycle genes. Ultimately, while our results also showed that complex carbon sources were likely being metabolized at the time of sampling (**Supplemental Results**), it is likely that phasins, carbon storage granules, and the glyoxylate shunt are critical components for MSC-1 survival during desiccation conditions.

To support our omics-based inferences, we measured respiration rates and protein content in each of our samples for each treatment (**Fig. S3**). In agreement with our carbon conservation hypotheses, soil analog samples were producing significantly less CO_2_ than broth samples (p ≤ 0.05), yet still contained about the same level of measured proteins (p > 0.05). This suggests a significantly higher carbon use efficiency (CUE) in the structure conditions than in liquid (p ≤ 0.05) and supports the idea that carbon conservation via the glyoxylate shunt is enabling *Ensifer* (and likely others) to retain their protein levels under conditions of low respiration. Our respiration measurements and metatranscriptomic expression analyses highlight that the ability of *Ensifer* to survive with minimal metabolic handoffs from *Streptomyces* is possibly due to its ability to use PHA as a carbon source and the glyoxylate shunt as a mechanism to conserve that carbon.

### Lowest moisture levels in soil analog demonstrate an increase in metabolisms related to chitin degradation and respiratory processes

While the environmental matrix had the most significant treatment effect, our moisture gradients showed significant differences in overall community expression (**Fig. S1**, **Fig. S4**). Across moisture treatments, isocitrate lyase was the functionally annotated gene with the highest expression and had significantly lower expression in low moisture compared to high moisture (base mean = 53,889, log_2_foldchange = -28.23). This is likely due to a decrease in flux through the glyoxylate shunt, with more carbon being funneled through the canonical TCA cycle, possibly due to more oxygen. In addition, over a quarter of the genes that were more highly expressed in 100% soil analog versus 25% soil analog were transporters (avg. base Mean = 112, avg. log_2_fold change = 24.2), and 62% of transporter expression was assigned to *Ensifer* (**Fig. S5**, **Table S2**). Most of the transporter genes expressed by *Ensifer* were branched-chain amino acid transporters suggesting that in high-moisture conditions, N is likely acquired from free peptides or necromass. However, we detected differential transcript abundance from multiple organisms (**Table S4**) for non-specific generalist transporters like PTS (phosphoenolpyruvate transport systems) and glucose transporters (gtsA), which likely contributed to the acquisition of GlcNAc and glucose.

Under low-moisture conditions, we detected an increase in the expression of GH18 chitinases by *Streptomyces*. While *Streptomyces* GH18 expression was significantly lower in the soil analog 100% incubations relative to liquid culture (p ≤ 0.05), 3 of the 4 *Streptomyces-*encoded GH18s are significantly more highly expressed in the lowest moisture treatment relative to the high moisture treatment. In support of microbial communities shifting towards respiratory metabolisms under drier conditions, we observed *Sphingopyxis* to display higher expression of respiratory metabolisms related to glycolysis (acetyl-CoA synthetase [K01895] under lower moisture, which could explain its increased abundance in the community (as measured by metagenomics). Further, our respiration measurements showed significantly higher respiration in the lowest moisture (p ≤ 0.05) (**Fig. S7**). Our proxy for carbon use efficiency (CUE) also showed a lower CUE in lower moisture, although this difference was not significant (p > 0.05). Additionally, overall metatranscriptomic activity is significantly higher in the soil analog 10% than in our fully saturated soil analog 100% (t = 3.3, p ≤ 0.05). Together, these results suggest that a decrease in moisture, and therefore hydraulic connectivity, likely establishes a mix of niches where A) oxygen and carbon are available (i.e., conducive to metabolisms like chitin degradation and canonical TCA cycle), and B) oxygen or carbon are not available (i.e., conductive to metabolisms like glyoxylate shunt, ethanol fermentation, and sulfur assimilation).

### Metabolite informed metatranscriptomics suggest high expression of osmolytes and possible biofilm genes are important bacterial responses to stress in our consortia

To understand how the different microbial expression patterns were influencing the metabolite profiles in our incubations, we collected metabolite data paired with our metatranscriptomes (**Fig. S8, Table S5**). We detected and annotated 138 unique known metabolites with nearly a quarter of them being differentially abundant across our treatments. Static metabolites across treatments consisted of diverse range of molecules, though sucrose was the only sugar detected. Other metabolites included amino acids, organic acids, and aromatic compounds. Differentially abundant metabolites included compounds having roles in critical biological mechanisms such as stress response (ectoine and trimethylamine n-oxide) (47) signaling (indole-3-acetic acid) (48), inhibition of biofilm formation (indole-3-carbaldehyde and undecanedioic acid) (49), promotion of biofilm formation (N-acetylputrescine) (50), and biofilm modification (hypoxanthine) (51).

Observed gene expression patterns explained the difference for 23 of 24 metabolites with significant differential abundance (p ≤ 0.05). Amongst these metabolites were ectoine and N-acetylputrescine (**Fig. 5**). Sparse partial least squares (sPLS) results showed that *Ensifer* had the highest Variable Importance in the Projection (VIP) index with regards to the predicted and observed measurements of ectoine and N-acetylputrescine. However, given that *Ensifer* was expressing genes at such a high level in the structured treatment samples, we constrained our sPLS results by examining whether *Ensifer* encoded and expressed the genetic machinery necessary to produce the predicted metabolites. The overall expression of genes that contribute to the production of ectoine and N-acetylputrescine positively correlated with the observed metabolite concentrations (**Fig. 5C**, **Fig. 5F**), suggesting that *Ensifer* is likely the main driver of these two metabolite concentrations, which can be important in processes like stress response (ectoine) and biofilm production (N-acetylputrescine).

## Discussion

### Phenotypes expressed in structured habitats differ from liquid cultures

Soil factors, including moisture and structure, are important controllers of microbial phenotypes under field conditions, but not often incorporated wholistically in mechanistic lab experiments. Here, we used our model soil consortia (MSC) resulting from a chitin-amended soil enrichment (15) to understand the constraints that moisture and the environmental matrix have on soil organic polymer decomposition (e.g., chitin). We provide the first available MAG representatives of the previously identified MSC-1, with all but 5 of our 29 MAGs being high-quality, near-complete genomes from long-read metagenomic sequencing. Further, using this tractable consortium, we show that overall community expression was significantly different when MSC-1 communities were cultivated in liquid versus soil analog (**Fig. 2**, **Fig. S1**), and that microbial respiration and biomass vary significantly between these conditions (**Fig. S3**, **Fig. S7**).

Differences in porosity and connectivity due to varying moisture levels in a structured growth matrix like soil can change redox conditions and SOM depletion rates (52). Reflecting this paradigm, *Streptomyces* and *Dyadobacter* showed a significant reduction in transcript abundance when grown in soil analog compared to liquid (**Fig. 2**). Previous studies observed that *Streptomyces* is central to chitin degradation in both a reduced complexity community (53) as well as a variety of ecosystems (54, 55). In agreement with prior experiments (53), the most highly expressed chitinase was encoded by *Streptomyces*. In fact, *Streptomyces* chitinase expression is 18-fold lower in 100% soil analog compared to liquid culture. We confirmed that all *Streptomyces* GH18 chitinases were extracellular enzymes, thus one possible explanation for these high expression patterns in liquid culture is that chitinases (and the resulting degradation products) diffuse away in a fully liquid system. As a result, it is possible that *Streptomyces* is being “cheated” out of chitin degradation products by microbial populations that do not contribute to chitin backbone decomposition. Community “cheaters” are more prevalent in well mixed systems that do not have a physical environmental matrix (56). In support of this, we saw that *Dyadobacter* accounted for 33% of the chitin byproduct degradation (e.g., chitobiose, GlcNAc-6P, GlcN-6p) in liquid culture, while contributing none of chitinase activity. Additionally, contrary to *Streptomyces* chitinase in the broth treatment, the most expressed chitinase (GH19) in the soil analog treatments was predicted to be a membrane-bound enzyme from *Ensifer*. As such, it is likely that by being membrane-bound, the chitinase yields higher concentration of privatized local chitin degradation products (i.e., chitooligomers and n-acetylglucosamine) to allow for species-specific growth, as opposed to widespread community sharing, a concept that has been previously explored (57).

### Motility, quorum sensing, and biofilm production in soil analog influence community chitin degradation

Another possible reason for the reduction of chitinase expression in soil analog is that *Streptomyces* are filamentous, non-motile bacteria, and are one of thirteen bacteria in this consortium that do not have a flagellar mechanism, along with *Dyadobacter* (**Supplemental Results**). Recent work found a positive relationship between flagellar motility mechanisms and cellular access to carbon (58), which supports the idea that the decrease in expression of *Streptomyces* (and other non-motile organisms) is possibly due to the inability to access substrates in soil analog incubations. Other work has also suggested a relationship between the ability of a cell to form flagella and produce biofilms (59), which may explain why *Ensifer* is so abundant and highly expressed in the structured incubations. Moreover, decreases in moisture content can lead to further isolation and differences in nutrient availability, and these dynamics are significant drivers of microbial processes across soil pores sizes (60), and varying hydrologic connectivity (61). Although we had sparse metaproteomic data, *Ensifer* encodes for 3 of the 5 most expressed proteins detected in the soil analog treatments which correspond to flagellin-related genes, and these genes have a 1.7-fold increase in soil analog compared to liquid culture, which suggest its importance for survival.

The role that spatial structure can have on a microbial community can also go beyond substrate acquisition. A recent publication implemented an individual-based spatial simulation and found evidence that spatial segregation was induced by competition avoidance (62). Mattei and Arenas propose that this competition avoidance is an important mechanism for the coexistence of microbial populations in communities. In addition to the proposed idea of metabolic “cheaters” and competition for substrates, it is likely that in structured habitats a lack of avoidance strategies (e.g., motility) decreases the competitiveness of certain non-motile organisms like *Dyadobacter* and *Streptomyces.* Ultimately, our ability to generate a reduced complexity soil structure analog that captures these critical ecological dynamics at low production cost could enable the constraining of computational models to more correctly predict microbial community behavior in natural soil settings.

The degradation of chitin (and other complex carbon types) in heterogeneous environments is referred to as a “public goods dilemma” that can be solved by generating biofilms that confine goods to producers (63, 64). Previous experiments have shown *Ensifer* can generate biofilms (65). Further, ectoine (66) and putrescine (50) are metabolites that are commonly associated with osmotic stress and biofilm production, respectively. Since our sPLS analyses link the metabolites for these processes to *Ensifer* gene expression, it is likely that *Ensifer* alters community metabolism under soil analog stress-inducing conditions which will ultimately influence oxygen gradients, substrate availability, and metabolic cross-feeding (67). Moving forward, being able to localize the degradation of chitin within a soil analog incubation (i.e., whether it occurs within a biofilm and in which layer) with paired biogeochemical measurements would help further constrain the ability of microbial communities to degrade chitin.

### Microbial stress and carbon conservation responses are defining factors of overall microbial community expression patterns in soil structure analog

How microorganisms allocate carbon (68) and how bioavailability of carbon is influenced by moisture (69) are critical components of ecosystem-level dynamics. As such, we set out to understand how carbon allocation was changing in response to moisture and environmental matrix across our treatments. Our results showed that while respiration (i.e., CO_2_ release) was highest in the liquid culture and lowest in 100% soil analog, biomass (i.e., total protein detected) remained consistent across treatments, suggesting metabolic adaptation. The glyoxylate shunt plays a key role in microbial adaptation to anoxia, carbon limitation, and desiccation (70–73). Throughout both the structured vs liquid and high-moisture versus low-moisture differential expression analyses, isocitrate lyase (K01637) is the highest differentially expressed, functionally annotated gene (**Fig. 4**). In agreement with a previous study that used qRT-PCR, isocitrate lyase, and not malate synthase, was the only glyoxylate shunt gene that was more highly expressed (72), which we hypothesize is due to regulatory signals. We also detected a decrease (but not an absence) in isocitrate lyase between the 100% moisture soil analog and 25% moisture soil analog, suggesting a bet-hedging mechanism employed to survive which decreases (but does not shut down) canonical respiration metabolism.

Microorganisms are known to adapt to environmental stressors to survive under a myriad of conditions. One of these survival strategies can be the production of polyhydroxyalkanoate (PHA) carbon storage molecules, which can store carbon during times of optimal substrate availability and tap into those resources under carbon-limited, stressful conditions (74–76). The most expressed functionally annotated gene in our entire dataset belongs to *Ensifer adhaerens* and is annotated as a phasin gene. Phasins are surface-binding proteins for the PHA granules. Previous literature states that *Sinorhizobium*, a genus very phylogenetically close to *Ensifer*, can use its isocitrate lyase gene alongside PHA, specifically poly-β-hydroxybutyrate (PHB), from its carbon storage granules to survive in carbon limited conditions (77). *Sinorhizobium* can also perform bet-hedging between high and low PHB-producing progeny, which enhances geometric mean fitness by producing daughter cells suited to both short-term and long-term starvation (78). Moving forward, pairing multi-omic measurements with visualization of granules will help further untangle the underpinnings of chitin degradation in these soil-relevant conditions.

### Reduced complexity consortia grown in soil analog sheds light on microbial priming dynamics for complex carbon degradation

Given how widespread and highly expressed phasins were in our samples, we hypothesized that PHA granules might play a priming role for the overall ability of MSC-1 organisms to use chitin as a carbon source. Our reduced complexity community was cultured in natural soil extracts which contained a myriad of different possible carbon sources besides chitin including glucose and sucrose (**Table S5**). Additionally, our incubations were amended with 600ppm of glucose to stimulate microbial growth in addition to the 1000ppm chitin. Respiration measurements from a preliminary experiment showed rapid growth patterns within 24 hours, at which point we hypothesize the cells were primed to produce enzymatic machinery for chitin degradation (**Fig. S9**). We also hypothesize that during this high-carbon earlier-stage, microbes are also producing carbon storage granules like PHA for future use.

These priming and growth constraint dynamics are well studied in soil and river hyporheic ecosystems (79–81). Specifically in soils, researchers have shown that glucose causes significant shifts in soil chemistry and antibiotic inhibition in *Streptomyces* (82). As such, we propose that for MSC-1, the overall ability to degrade chitin depends on the availability of simpler carbon substrates that can kickstart microbial metabolism. Specifically, we hypothesize that *Streptomyces* can only achieve high levels of chitin degradation in liquid systems that maximize diffusion of priming substrates and metabolic cross-feeding, or within biofilms that can facilitate the organization and transfer of public goods. Consequently, we suspect that the observed patterns of decreased chitin degradation in the soil analog condition are a combination of lack of motility, priming substrates, and suitable redox conditions, which together activate carbon conservation strategies that are more suited to those conditions (i.e., *Ensifer* glyoxylate shunt). Given the reproducibility of our soil analog incubations, we believe that our model consortia is a great starting point for identifying metabolic and environmental control points dictating chitin decomposition.

Recent work demonstrated a drought induced trade-off between stress tolerance traits and resource acquisition traits (83). Induction of such strategies ultimately should alter carbon flux and energy expenditure for these organisms and would likely alter SOM decomposition rate and CUE, or the proportion of growth over carbon intake. Here, we find support for these findings using a reduced complexity model consortium and present evidence that habitat structure, and to a lesser degree moisture, induce altered community metaphenome resulting from A) a metabolic shift to stress tolerance traits (e.g., increased production of ectoine and n-acetylputrescine), B) a decrease in resource acquisition traits (chitinase expression), and C) yield optimization via an increase in enzymes involved in carbon conservation pathways including glyoxylate shunt and PHA utilization. The complexity of these phenotypic responses are also key reminders of the emergent properties that arise out of microbial community interactions which are difficult to track in their entirety. Even at a scale used here of approximately 29 different organisms, phenotypic expression was more than just the target chemical pathways that these communities were enriched for (chitin degradation), and instead are a complex network of interactions related to redox conditions (i.e., oxygenation, porosity, connectivity), metabolic handoffs (i.e., community cheaters and biofilm production), and fluctuating carbon utilization metabolisms (i.e., glyoxylate shunt, glycolysis, chitin metabolism). Ultimately, given the effectiveness of the soil analog-grown model soil consortia in interrogating microbial metabolism in heterogeneous conditions, this system can be of use to the broader community by providing a tractable, higher-complexity alternative to microbial studies in liquid cultures.

## Data Availability

All code generated for this study is publicly available on GitHub (https://github.com/jrr-microbio/MSC1_SSA_incubations_moisture) via an R markdown document. Raw reads, trimmed reads, and assemblies are available on the JGI data portal under project IDs: 1404644-1404648 (long-read metagenomes), 1404665-1404669 (short-read metagenomes) 1404747-1404771 (metatranscriptomes). MAG fasta files are hosted for reviewers on Zenodo: https://zenodo.org/records/13834436 and will be permanently hosted after review in the PNNL DataHub Search | Datahub (pnnl.gov).

## Acknowledgements

This program is supported by the U. S. Department of Energy, Office of Science, through the Genomic Science Program, Office of Biological and Environmental Research, under FWP 70880. Protein and metabolite analysis by LC-MS was performed in the William R. Wiley Environmental Molecular Sciences Laboratory (EMSL), a national scientific user facility sponsored by Office of Biological and Environmental Research and located at PNNL under EMSL project 60461: Prediction of response of microbial interaction networks cycling carbon to changing moisture conditions. PNNL is a multi-program national laboratory operated by Battelle for the DOE under Contract DE-AC05-76RLO 1830. Metagenome and metatranscriptomes were generated at the DOE Joint Genome Institute on Project 508623 under Contract No. DE-AC02-05CH11231. Thanks to Patricia Miller who assisted metabolite sample preparation, Josie Eder who analyzed the samples by LC-MS, and Isabella Yang who assisted with metabolite identification.

---

**Supplementary figure 1:**
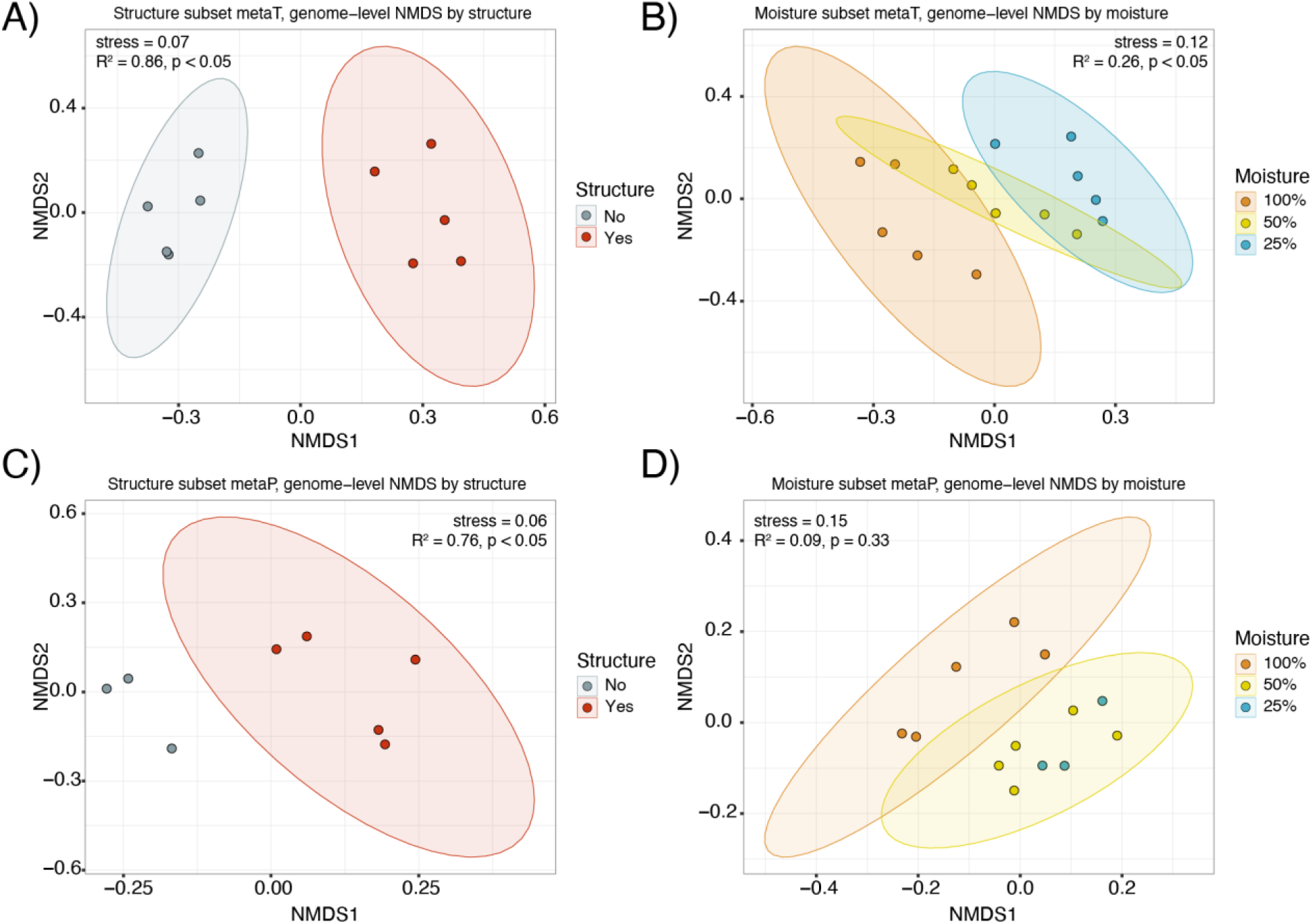
Structure and moisture influence community expression in metatranscriptomes and metaproteomes. A-B) NMDS shows structure and moisture drive differences in community expression by metatranscriptomes. B-C) NMDS shows structure and moisture drive differences in community expression by metaproteomes. There were not enough samples to draw a confidence ellipse for the no-structure ellipse in (C), nor the 25% moisture in (D).

**Supplementary figure 2.**
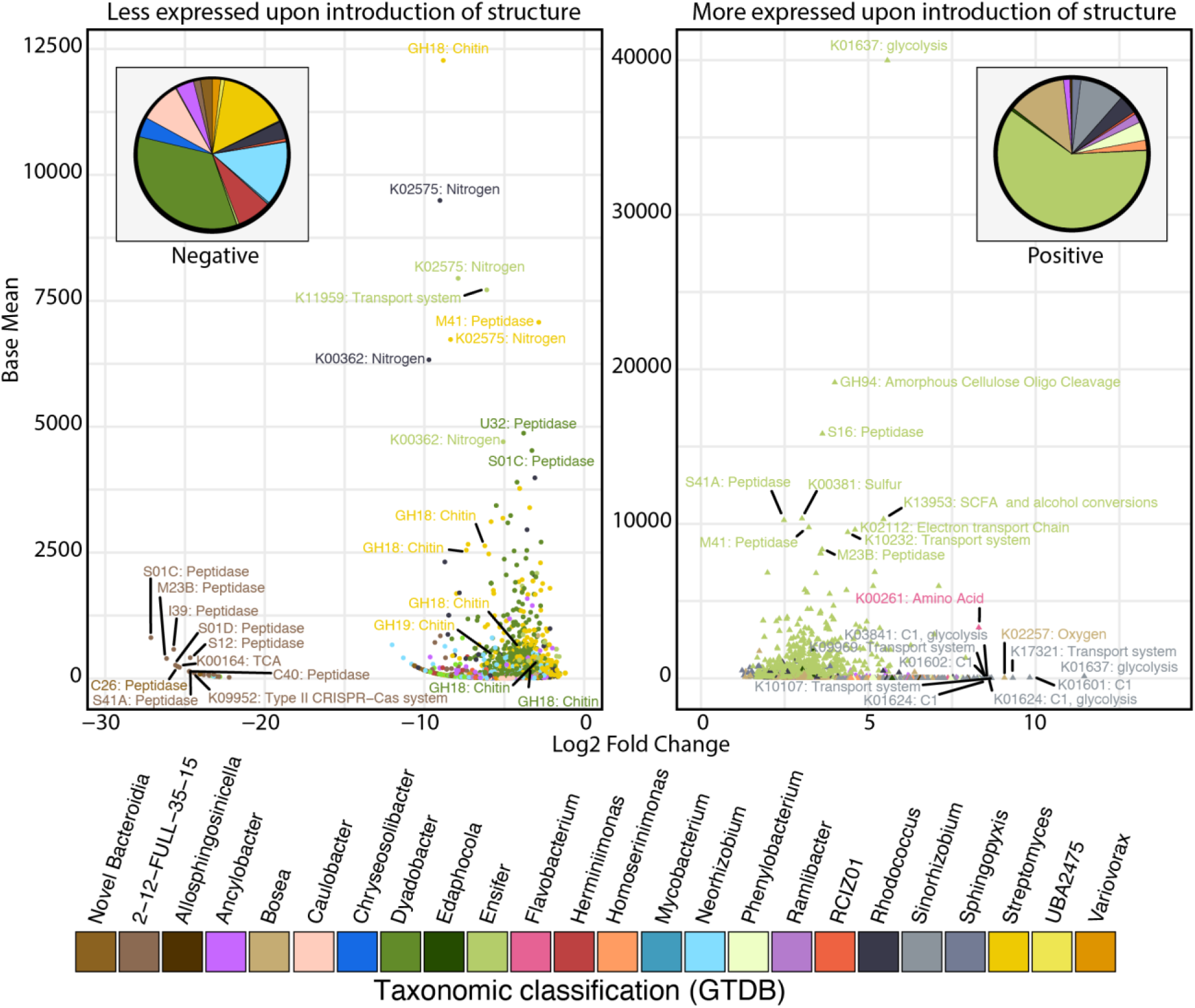
Differentially expressed genes across structure treatments highlight decrease in chitin degradation and increase in alternative microbial lifestyles. Dot plots show significantly differentially expressed genes (DEGs) across soil analog treatments. Whether those differentially expressed genes are lower (left plot) or higher (right plot) is denoted by shapes. The x-axis shows the log_2_ fold change of each gene, and the y-axis denotes the base mean expression of each DEG. Colors denote the taxonomic classification of each of the genes. Pie charts denote the overall distribution of organisms that had higher or lower expression of genes. The functional annotation is shown for a subset of genes.

**Supplementary figure 3:**
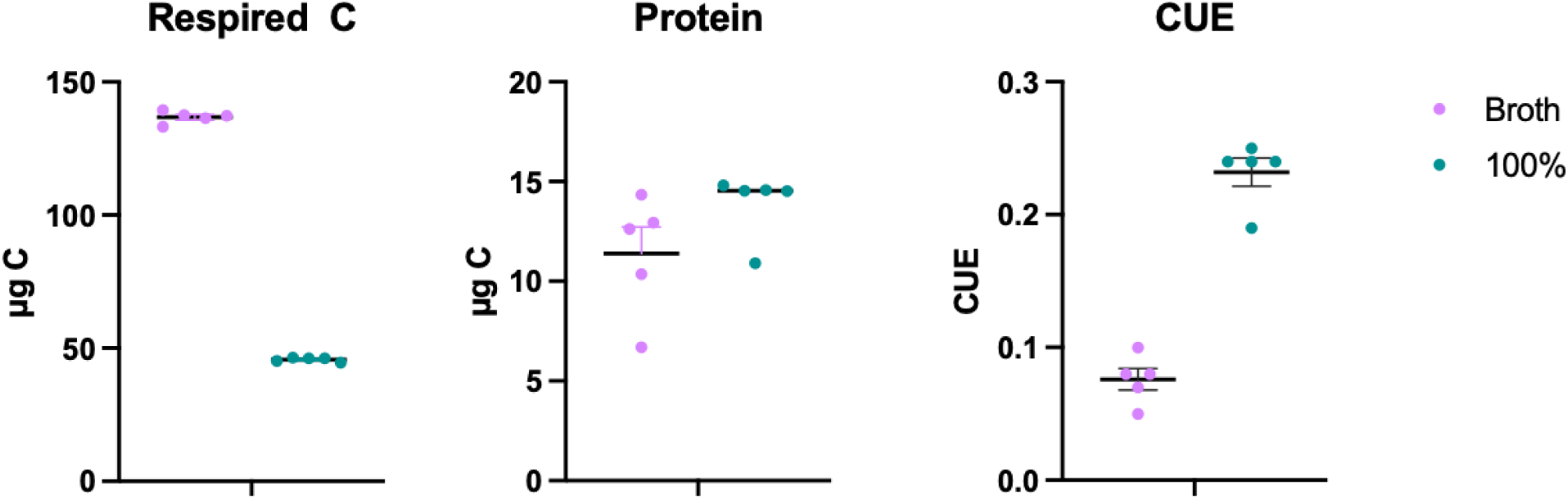
Carbon Use Efficiency proxy (CUE). Carbon allocation to cumulative respiration and cellular protein, as proxy for biomass, was quantified at 96 h of incubation and used to calculate the apparent carbon use efficiency (CUE) for broth and 100% moisture saturation SSA.

**Supplementary figure 4.**
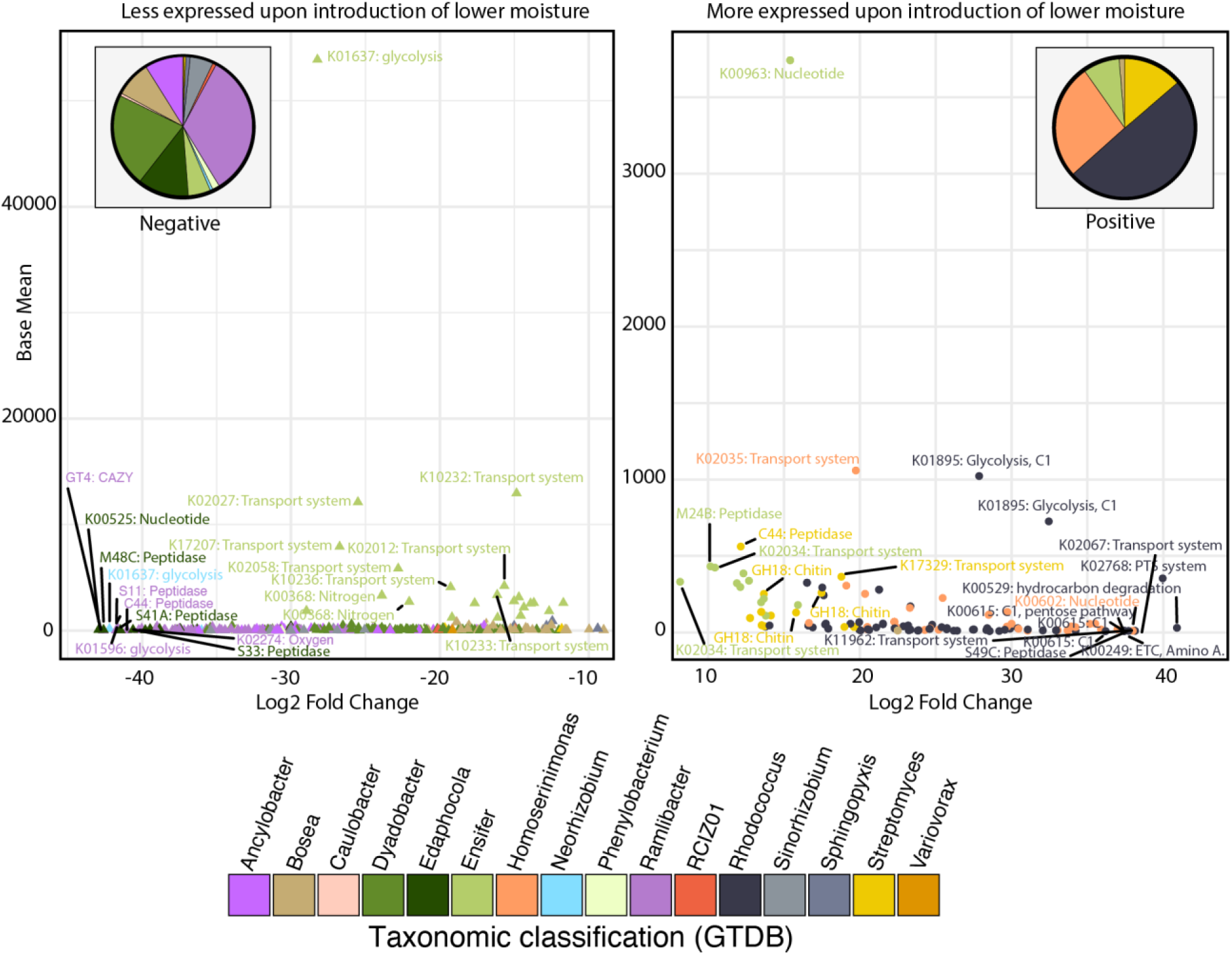
Differentially expressed genes across high and low moisture treatments highlight slight increase in chitin degradation. Dot plots show significantly differentially expressed genes (DEGs) across structure analog moisture treatments. Whether those differentially expressed genes are lower (left plot) or higher (right plot) is denoted by shapes. The x-axis shows the log_2_ fold change of each gene, and the y-axis denotes the base mean expression of each DEG. Colors denote the taxonomic classification of each of the genes. Pie charts denote the overall distribution of organisms that had higher or lower expression of genes. The functional annotation is shown for a subset of genes.

**Supplementary figure 5.**
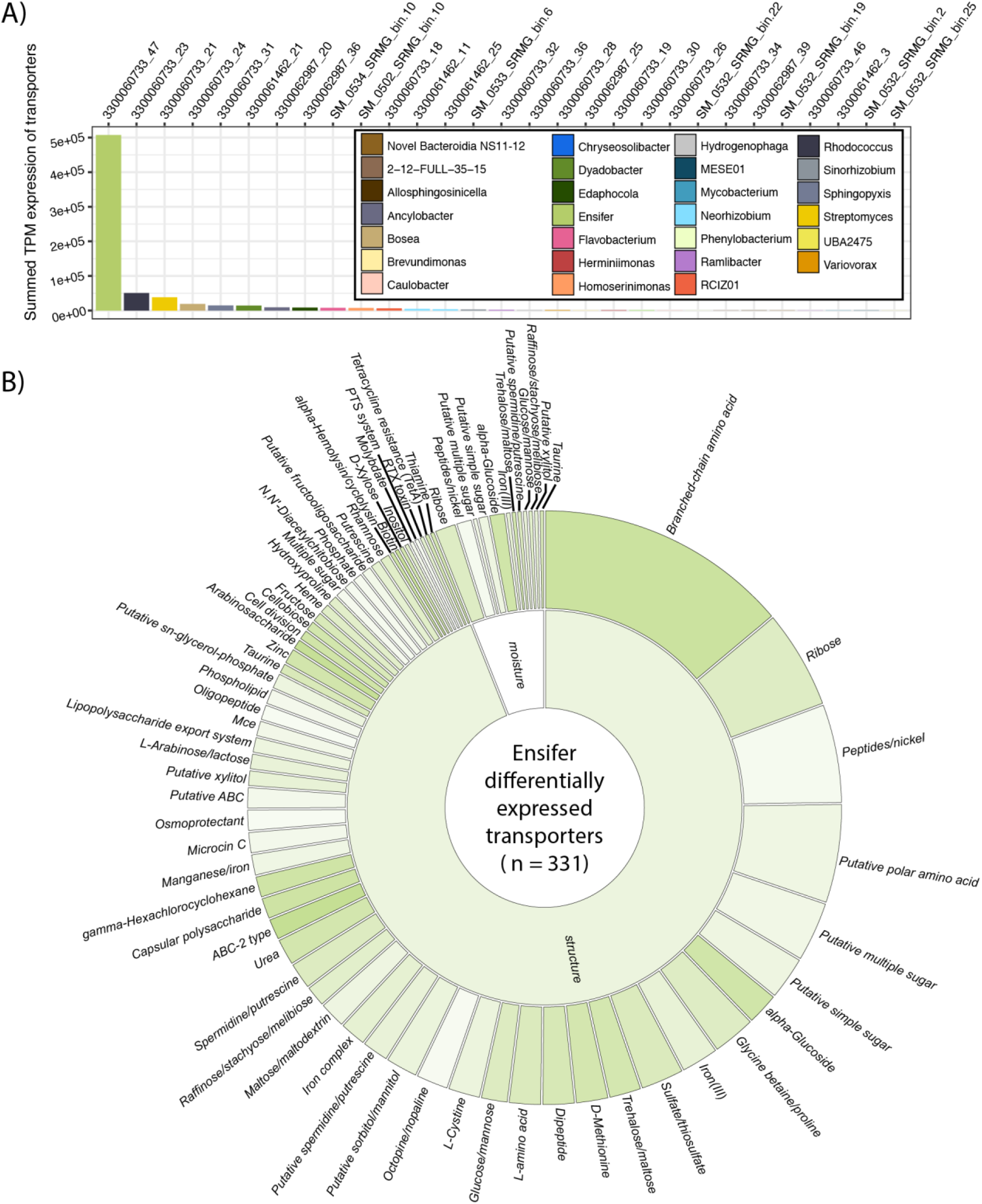
Stress response and scavenging metabolisms upon differential moisture regimes. **A)** Boxplots show the overall summed TPM expression of transporter genes per genera. **B)** Sunburst diagram highlights the functional categorization of transporters that are differentially expressed across treatments by *Ensifer sp*.

**Supplementary figure 6:**
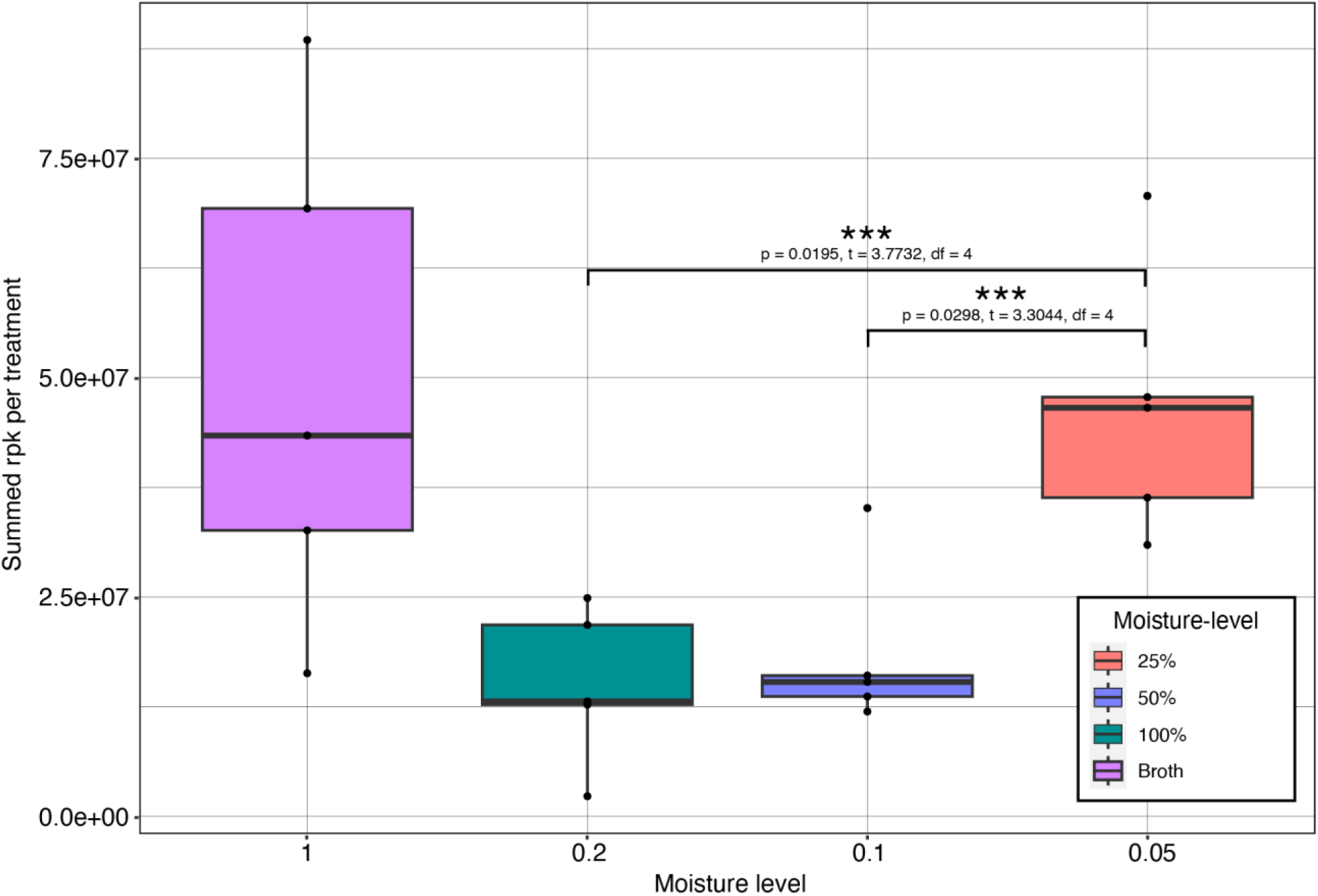
Overall rpk normalized expression values of full communities across treatments. Bar graphs denote the total rpk normalized expression of all MSC1 community members. Significance is shown by horizontal bars on top of bar plots (when applicable). Colors denote the different structure / moisture treatments.

**Supplementary figure 7.**
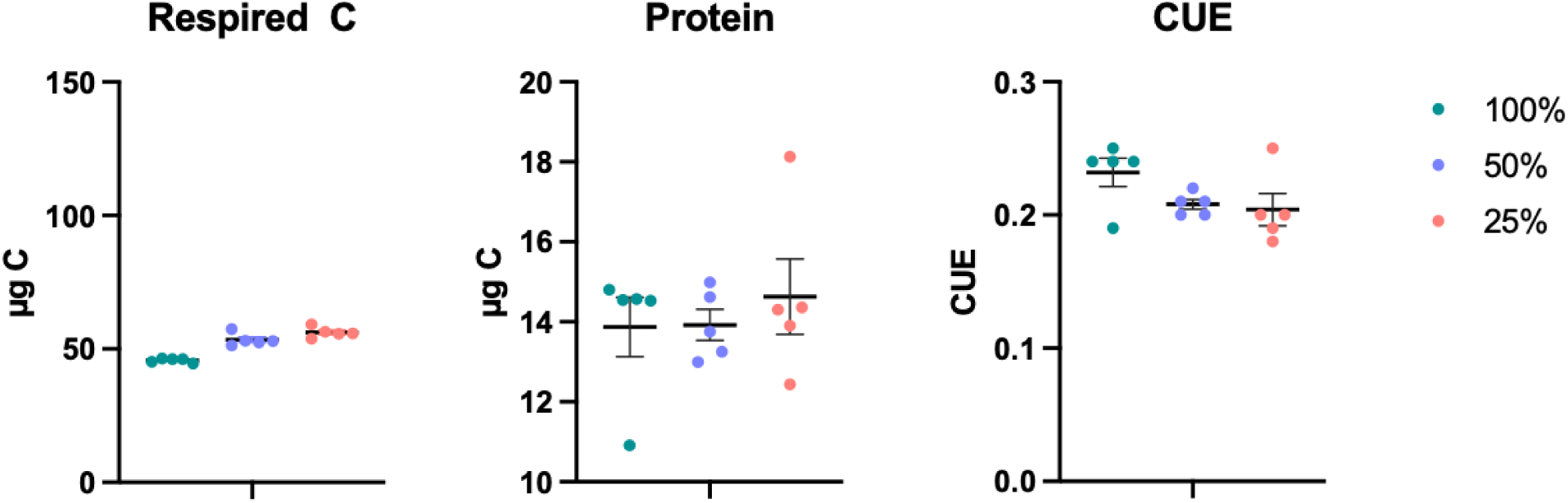
Carbon use efficiency plots across moisture. Carbon allocation to cumulative respiration and cellular protein, as proxy for biomass, was quantified and used to calculate the apparent CUE for 100%, 50%, and 25% moisture saturation SSA.

**Supplementary Figure 8:**
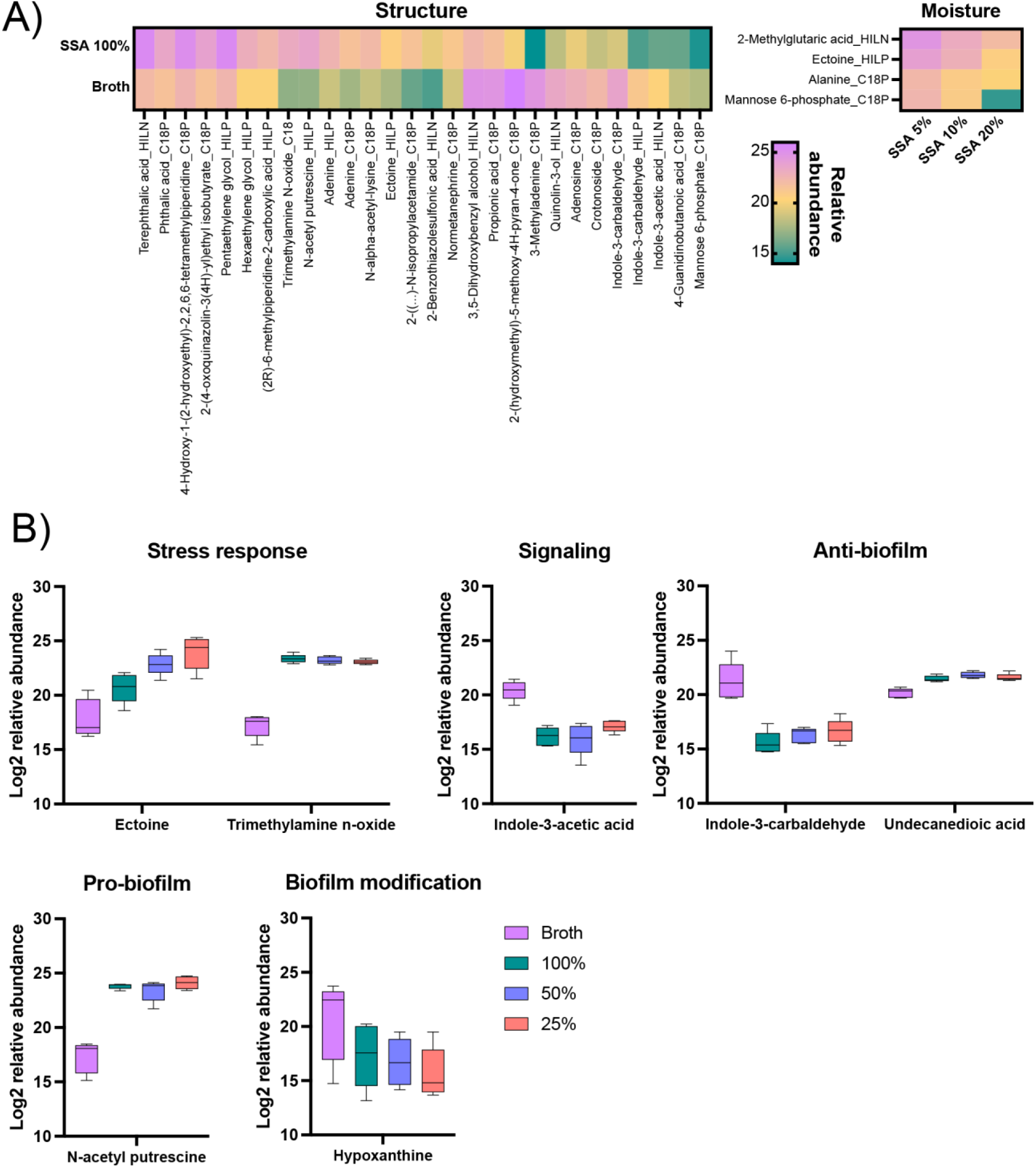
Metabolite concentrations. **A)** Median normalized relative abundance values were log_2_ transformed and the heat hap depicts the metabolites that had significantly different values (p ≤ 0.05) based on either structure (broth vs 100% SAA) or moisture (100%, 50%, or 25% SSA). **B)** Key extracellular metabolites of interest. Significant structure and or moisture effects on metabolite relative abundance with log_2_ fold change ≥ 2, p ≥ 0.05.

**Supplementary Figure 9:**
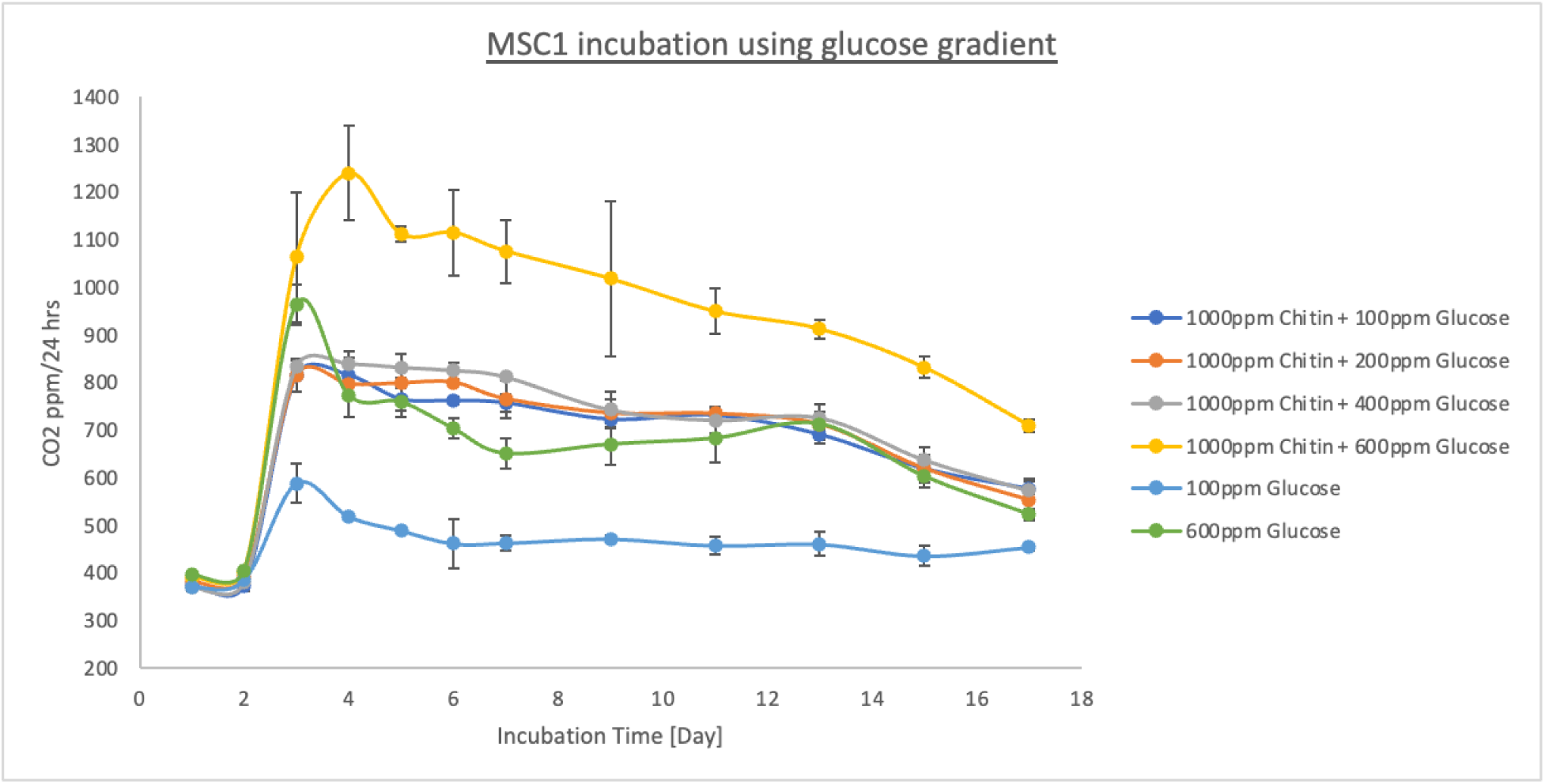
Respiration curves of glucose-amended chitin incubations. The y-axis denotes the total respiration that was measured within 24 hours of the x-axis denoted incubation time. Different line colors show different tests that were performed with variable levels of glucose and chitin.

**Supplementary figure 10.**
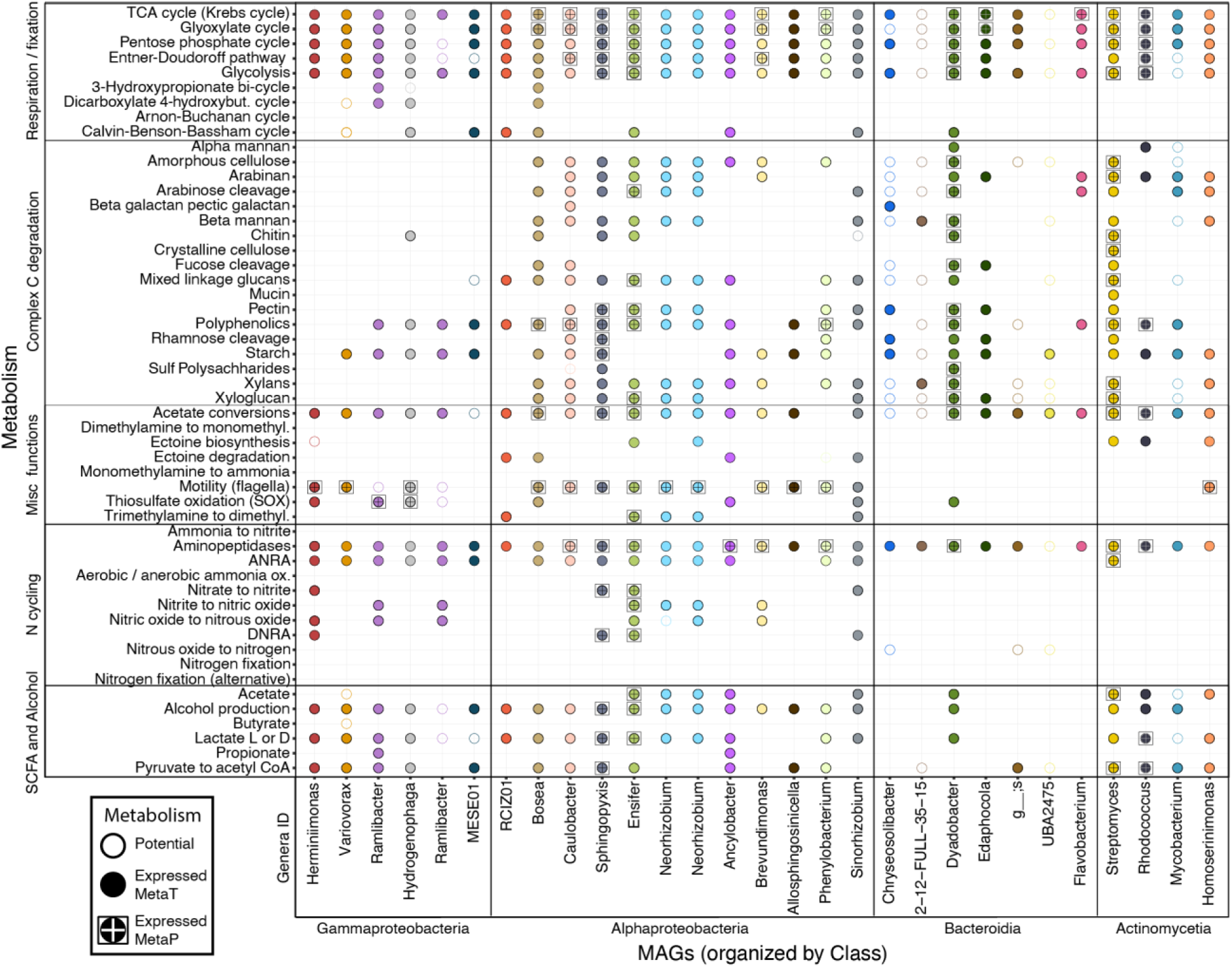
MSC-1 microbial communities are functionally redundant. Heatmap shows metabolisms for each genome. Empty circles denote metabolic potential. Filled circles denote expression detected by metatranscriptomics. A cross inside each circle and a bounding box surrounding it denotes that expression is further supported by metaproteomic data.

